# Automated quality control of next generation sequencing data using machine learning

**DOI:** 10.1101/768713

**Authors:** Steffen Albrecht, Miguel A. Andrade-Navarro, Jean-Fred Fontaine

## Abstract

Controlling quality of next generation sequencing (NGS) data files is a necessary but complex task. To address this problem, we statistically characterized common NGS quality features and developed a novel quality control procedure involving tree-based and deep learning classification algorithms. Predictive models, validated on internal data and external disease diagnostic datasets, are to some extent generalizable to data from unseen species. The derived statistical guidelines and predictive models represent a valuable resource for users of NGS data to better understand quality issues and perform automatic quality control. Our guidelines and software are available at the following URL: https://github.com/salbrec/seqQscorer.

## Background

Functional genomics based on next generation sequencing (NGS) technology is used to study regulatory elements in genomes of all types of species. It is widely used in biological and clinical applications thanks to a variety of existing complementarity assays that allow the investigation of, for example, gene expression quantification (RNA-seq), epigenetic modification and transcription factor occupancy (ChIP-seq), and open chromatin regions (e.g. DNAse-seq, MNAse-seq or ATAC-seq).

The analysis of NGS data requires a stepwise process handled by dedicated software tools. The first processing step is the quality control (QC) of the data. It is of crucial importance to filter out low quality data files that would have a negative impact on downstream analyses through addition of noise or systematic bias to the analyzed dataset [1]. When deriving differences between groups of samples, low-quality samples would increase the variance within a group and thus hamper the ability of statistical tests to find significant differences amongst them. In a clinical context, patient data of unnoticed low-quality can also lead to wrong diagnosis or ill-suited treatment. Filtering out or editing a small portion of sequencing reads within a file or applying more sophisticated bias mitigation methods may be detrimental to the downstream analysis [2], [3], or may not be enough to correct such a systematic error [4]. Common QC tools analyze the data files to derive numerous highly specific quality features for manual review. As the usefulness of many of these features was never demonstrated, a large majority of NGS scientists is still not confident about classifying a sequencing file by quality.

QC tools, used at the first step of an NGS pipeline, analyze raw sequencing data. The raw data is stored in FastQ files containing a set of short string of DNA sequences (e.g. 100 bases long) called reads, and related information such as a quality score for each base reflecting its sequencing error probability. The most popular QC tool in the NGS community is FastQC (http://www.bioinformatics.babraham.ac.uk/projects/fastqc/). It performs various analyses that could indicate problems such as position-dependent biases (“Per base sequence quality” analysis), sequencing adapter contamination (“Overrepresented sequences” analysis), or DNA over amplification (“Sequence duplication levels” analysis). Quality analyses based on the raw data can be complemented by analyses performed at later steps of the NGS data processing. An important subsequent step is the mapping of the reads to the reference genome if available. Dedicated software tools such as Bowtie2 output mapping statistics that could be used as indicators of quality [5]–[8]. The number of sequencing reads that map to a unique position or the number of reads that do not map in the reference genome are presumably very important quality features. In following data processing steps, related software tools are often assay specific [8]–[12]. Their results could still complement the tools mentioned above for different assays. Analyzing the genomic location of the reads to know if they map predominantly to expected functional elements or to know the distribution of the reads near gene transcription start sites (TSSs) are of special interest to ChIP-seq, DNAse-seq or ATAC-seq data for example. Although some tools offer reports that integrate results from several QC software [13], [14], the final QC decision still remains manual. This decision is complex given the multiplicity of quality features generated at different steps of the data processing, their expected dependency from experimental conditions (e.g. species, assays or treatments), and the lack of statistical studies that would recommend specific values that differentiate low- and high-quality data. Therefore, a system to aid NGS QC related decisions, making them automated and independent of human biases, is desirable.

Due to the coordinated efforts of many research groups, large repositories have been created that collect NGS data files in order to make them available to the scientific community. The scope of some repositories such as GEO and ArrayExpress is to share data with a minimal amount of annotation describing experimental conditions. Annotations are created in accordance with detailed guidelines [15] and ontologies [16]. Although they help to maintain a high standard of data annotation, they do not control the quality of the deposited data. Other specialized repositories, such as TCGA for cancer data, focus on high-quality data. Uniquely, the ENCODE repository [17], [18], specialized in functional genomics, provides access to a large number of high- and low-quality NGS files that were labelled either as released or revoked, respectively, based on a semi-automatic QC procedure. Although ENCODE guidelines were created to help NGS specialists to produce data of high quality, curators of their repository still manually decide the quality of the files after reviewing various quality features [19], [20]. The goal of this study is to improve NGS QC procedures by comparing these files and applying statistical methods and machine learning algorithms to derive useful statistics and classification models leveraging comprehensive quality features.

Although machine learning has been used to classify the quality of reads or single-nucleotide polymorphisms [21], we have not found such an application to full NGS files. We focused our work on RNA-seq, ChIP-seq and DNAse-seq data files of human and mouse samples. We were first interested in defining the scope of application and relevance of each individual feature by comparing data statistics from different species and assays. Then, we used machine learning methods to derive optimal and unbiased predictive models combining multiple features. After extensive validation of the models, we investigated their usefulness in different scenarios such as database curation and disease diagnosis. Finally, we released to the community our statistical guidelines and a software application to use the classification models on new datasets.

## Results

We have studied a large number of annotated files to characterize data quality and to create machine learning models able to automatically predict quality from the raw data. Our goal is to provide an alternative to manual quality control of NGS data files, which currently requires high-level expertise and highly depends on assays and experimental conditions and is prone to human biases.

### Workflow

Our study is based on 2,642human and mouse raw NGS data files labelled as high- or low-quality from the ENCODE data portal (Figure 1). From them, we extracted different quality features using software tools commonly used in the NGS field. Each tool represents a different stage or view of the NGS data analysis, providing features sets related to raw sequencing reads (RAW), mapping to a reference genome (MAP), genomic localizations of the reads (LOC), and spatial distribution of the reads near transcription start sites (TSS) (see Methods for details).

**Figure 1.**
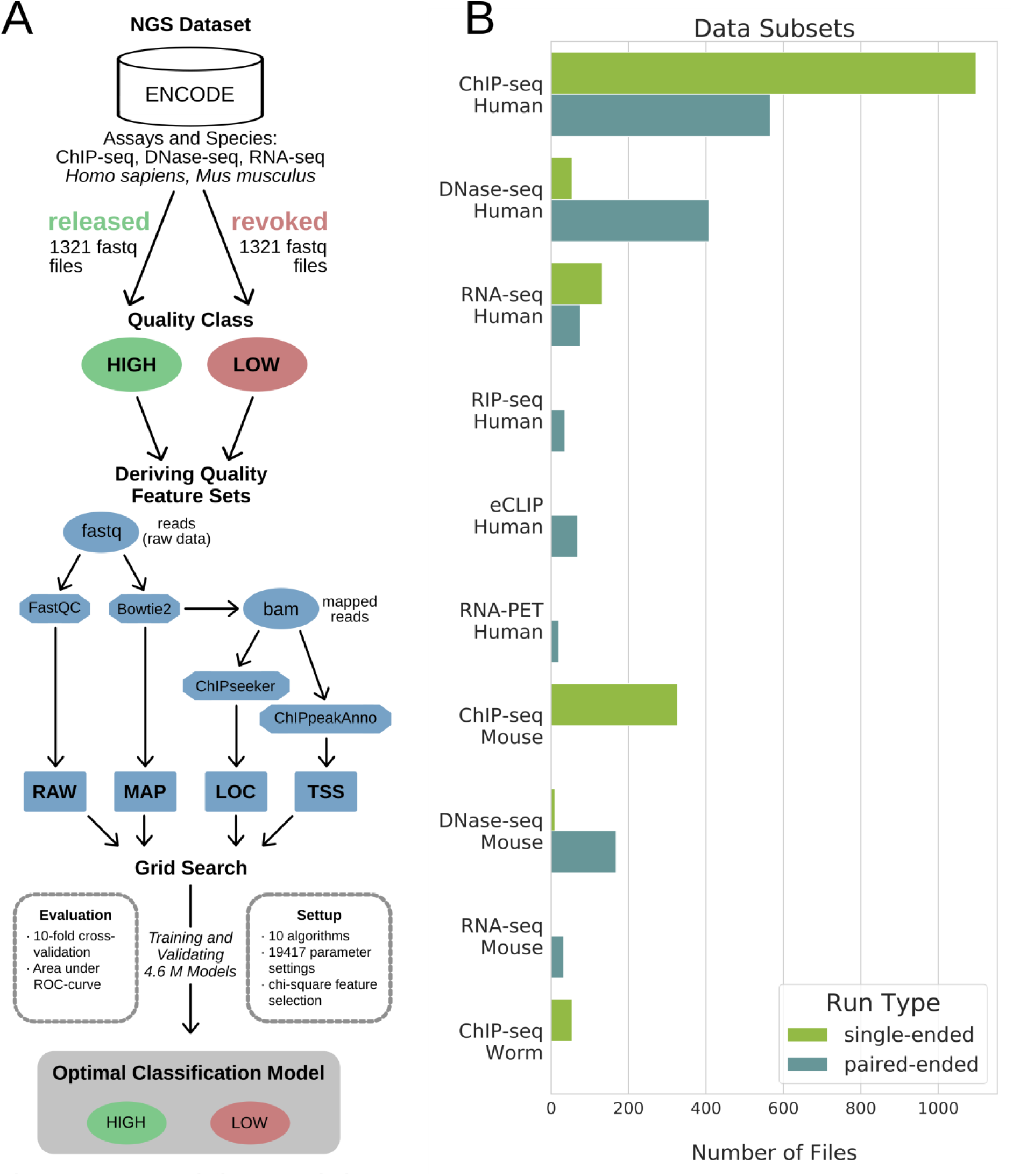
Workflow and dataset. **A** Workflow. Raw NGS data files were retrieved from the ENCODE data portal. Four quality feature sets were derived using standard bioinformatics tools (e.g. FastQC): raw data (RAW), genome mapping (MAP), genomic localization (LOC), and transcription start sites profile (TSS) feature sets. A grid search was used to derive optimal machine learning models by testing various parameter settings and algorithms. **B** Dataset. Bars show the number of files in each data subset (y-axis).

Our approach was first to derive statistical guidelines from the detailed study of individual quality features, and then to systematically benchmark 10 popular machine learning algorithms, such as multilayer perceptron or random forest (RF), to predict the quality of NGS data files based on combinations of quality features and various sets of parameters. We finally evaluated 4.6M predictive models covering different data subsets: either generic (including all species and/or all assays) or specialized in particular species and assays (e.g. human ChIP-seq or mouse DNase-seq).

### Quality prediction

#### One-feature quality predictions as baseline and guidelines

Before evaluating machine learning algorithms, we evaluated the predictive power of each quality feature independently (Figure 2A). This analysis provided us an overview of their performance across the data subsets and a baseline for the evaluation of the machine learning algorithms. We compared the results using the area under the receiver-operating-characteristic curve (auROC) that ranges from 0.5 to 1, from not predictive to completely predictive, respectively. The predictive power of the features strongly depends on the data subset, ranging from poorly predictive (e.g. genomic localization in “1^st^ exon” for human RNA-seq; auROC = 0.50) to highly predictive (e.g. “overall mapping” for mouse ChIP-seq files; auROC = 0.94).

**Figure 2.**
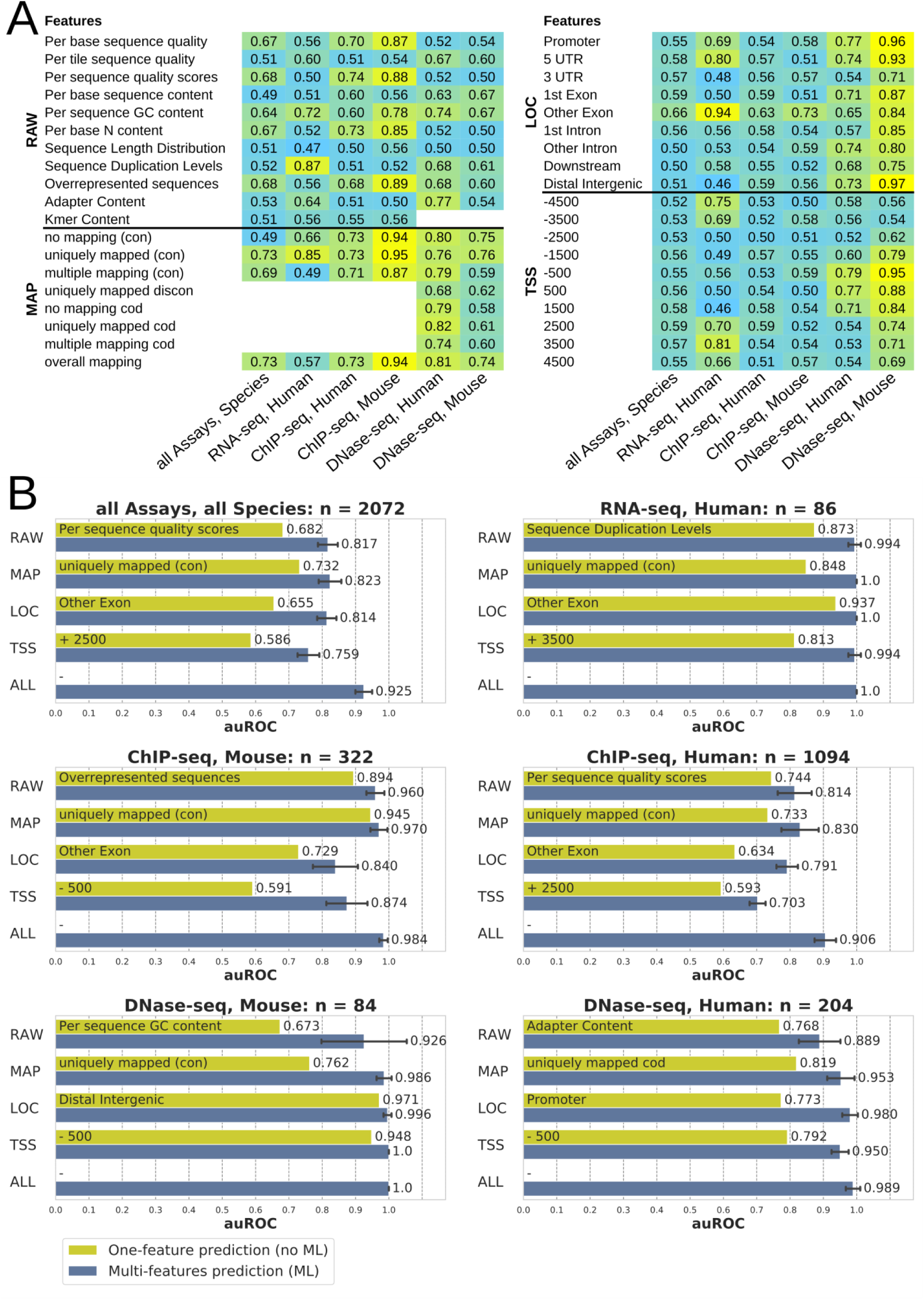
Predictive performance of quality features and machine learning (ML) **A** One-feature predictions. Predictive performances of each feature (no machine learning) as areas under ROC-curves (auROC) ranging from 0.5 (random predictions) to 1 (perfect predictions). **B** Multi-feature predictions. Predictive performances of optimal machine learning models trained using different feature sets (y-axis) outperform one-feature predictions. Error bars show standard deviations derived from 10-fold cross-validations within the grid search. Feature sets: RAW (raw data), MAP (genome mapping), LOC (genomic localization), TSS (transcription start sites profile), ALL (all features).

Some quality features are of broad interest because of their good performance across all data subsets, especially all MAP features and the following two RAW features: “Overrepresented sequences” and “Per sequence GC content” (auROC up to 0.89 and 0.78, respectively). Other quality features are less interesting because of their poor performance in all subsets (e.g. “Sequence Length Distribution”; or “TSS −2500”; auROC <= 0.62).

From this analysis, we have derived detailed statistical views that confirm the specificity to data subsets observed above (Figure S1). By providing an easy way to compare quality features of new data files to thousands of ENCODE files, these results can serve as statistical guidelines to NGS specialists.

#### Multi-feature quality predictions by machine learning

With the performance of each single quality feature as baseline, we evaluated the potential improvement of using combinations of features by machine learning algorithms to predict NGS data files quality. We evaluated machine learning models trained with the different data subsets for each feature set (Figure 2B). Models, tuned by a grid search that systematically explores performances over the parameter space, outperformed the baseline. Data files quality from each subset can be predicted with high performance (auROC > 0.9). Within each data subset, the different feature sets led to comparable results, although performances were more variable with RAW features in DNase-seq subsets, and LOC and TSS features sets were less performing for ChIP-seq subsets. This analysis confirms the expectation that sequencing reads from the ChIP-seq subset would not be highly biased towards specific genomic localizations or TSS relative positions different to RNA-seq, for instance, for which these two feature sets enable very high performing models (auROC=1 or 0.994, respectively). Results for all combinations of feature sets and other performance measures such as area under precision-recall curve, accuracy, or F1 are shown in supplementary (Figure S2 and Figure S5 A).

### Within-experiment analysis

The extent to which individual NGS researchers would benefit from using machine learning models in their QC procedure could be known by analyzing the results on replicate files produced within a same experiment. In our dataset derived from ENCODE, some experiments gather high- and low-quality files from different replicate and control samples. For the different species-assay combinations we extracted these experiments with the corresponding files (Figure 3A).

**Figure 3.**
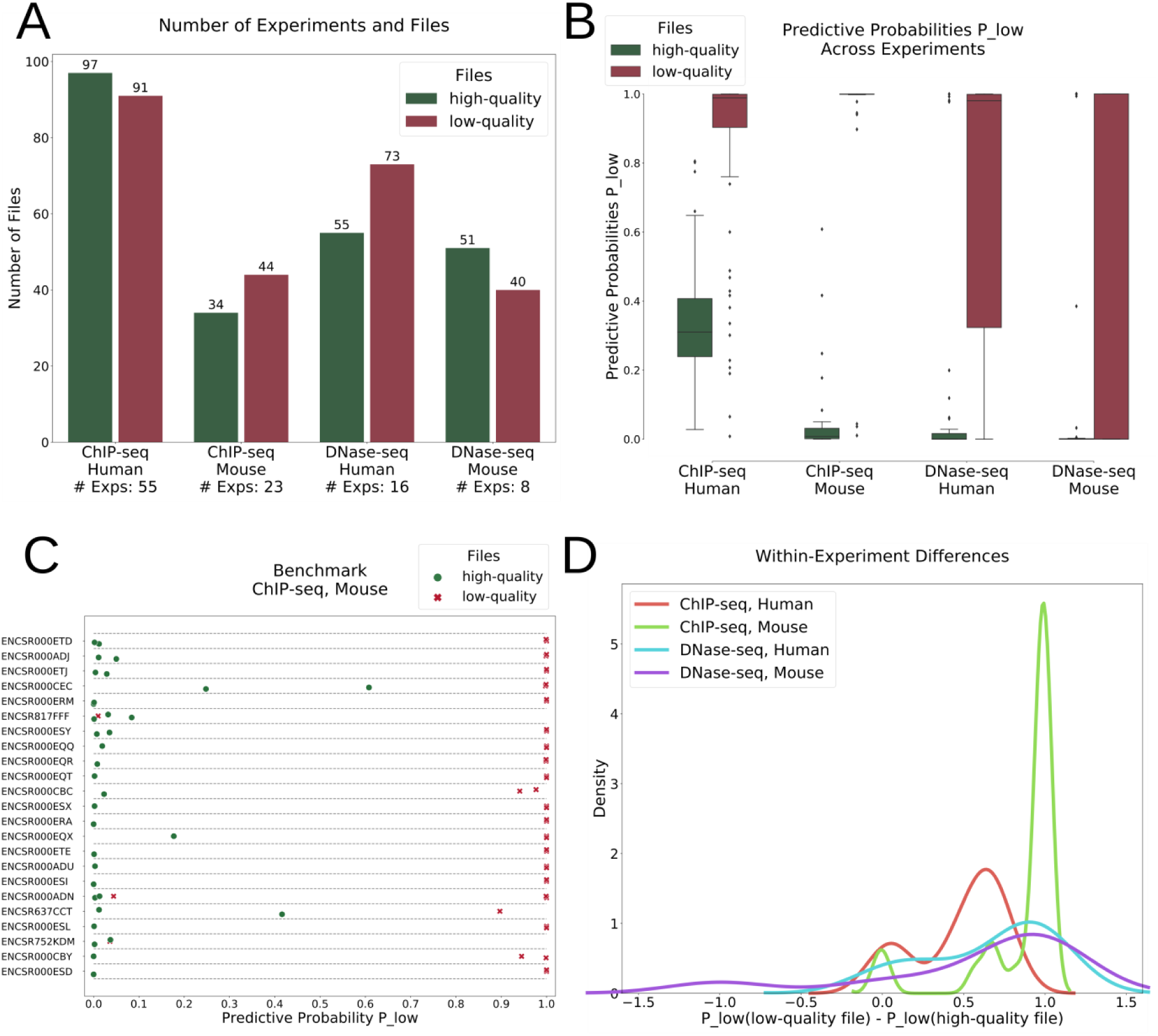
Within-experiment differences. **A** Numbers of selected experiments and files. Only experiments that contain released files together with revoked files were selected. **B** Predictive probabilities across experiments summarize the good predictive power in each data subset. **C** Benchmark of mouse ChIP-seq files in selected experiments illustrates the good predictive power of the models. P_low_: predictive probability to be of low-quality. **D** Density curves of within-experiment differences between predictive probabilities for low- and high-quality files. Average differences are large (range from 0.5 to 1).

Focusing on the probability of a file to be of low quality that was provided by the models to each file across the different experiments, our models were able to clearly differentiate which files would be considered high- or low-quality after manual QC (Figure 3B). Within each experiment, we could mostly observe a clear cut or a meaningful sorting by actual quality (Figure 3C and Figure S3), with an average difference between low- and high-quality files ranging from 0.5 to 1 (Figure 3D). From these observations, we conclude that the models were not biased towards some experiments. These results suggest that, early in the sequencing analysis pipeline, researchers can already define the potential of their data files to be considered of enough quality for publication and can accordingly take decisions that could save substantial amount of time and resources for further analyses, storage or manual reviews. For instance, out of 23 mouse ChIP-seq experiments in our dataset, 41 (53%) files could have been early identified as of low quality and not submitted, stored, processed and manually reviewed by the ENCODE database curators.

### Top machine learning algorithms and parameters

Out of our model tuning strategy, which systematically tested 10 different algorithms and numerous parameters sets, different algorithms could be found in models optimal for each data subset and/or feature set. Given the heterogeneity of the data composed of numerical and categorical values, and the moderate dataset size, we thought that decision-tree-based algorithms would be appropriate to the task. To test this hypothesis, we summarized the results across the data subsets and combinations of feature sets. As expected, decision-tree-based algorithms (random forest, gradient boosting, and XG boost) often performed better than others as well as multilayer perceptron, which is a deep learning classifier based on artificial neural networks (Figure 4A). In general, there were only minor differences between the top algorithms. Nevertheless, the choice of their parameters proved to be critical (Figure 4B). For example, the multilayer perceptron clearly benefited from the quasi-Newton method (lbfgs) to solve the weight optimization (>80% of the best models) but optimal sizes of the hidden layers were more dependent on other parameters. The choice of an automatic features selection could also be of importance, especially with gradient boosting algorithms (Figure 4C).

**Figure 4.**
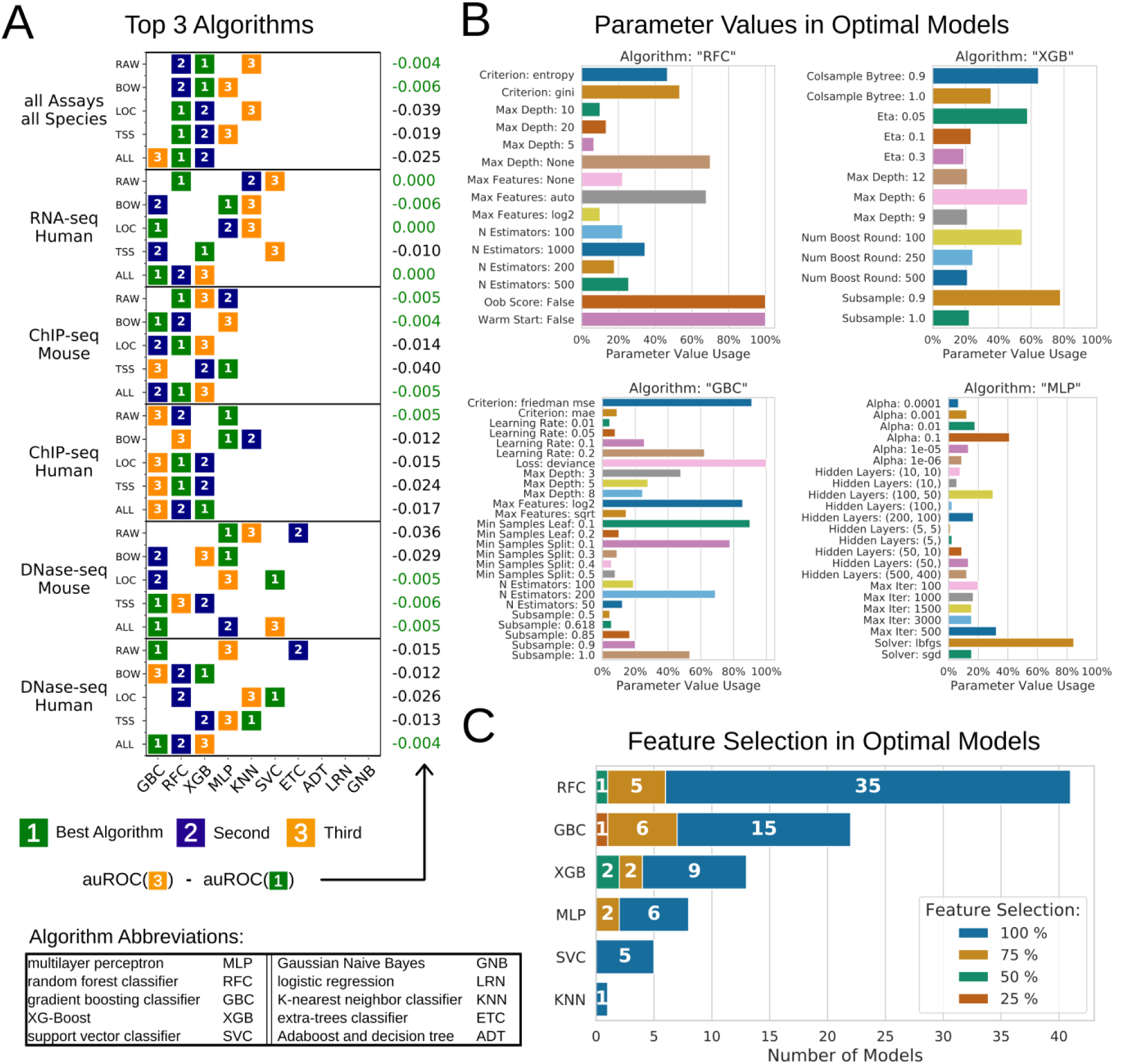
Best performing algorithms and parameters. **A** The top three performing algorithms for selected data subset-feature set combinations (colored boxes). Numbers on the right-hand side show the difference of the area under ROC-curve (auROC) between the very best and the third best algorithm. Feature sets: RAW (raw data), MAP (genome mapping), LOC (genomic localization), TSS (transcription start sites profile), ALL (all features). **B** Frequency of algorithmic parameter settings in 90 optimal models. For each algorithm, optimal models were created for each possible combination of feature set and data subset. **C** Frequency of chi-square-based feature selection settings in best performing models. For each possible combination of feature set and data subset, we compared optimal models derived by different algorithms and retained the best one and its algorithm.

### Cross-species generalization

A main limitation of this study is the availability of labelled data only for human and mouse. In order to know if the models were potentially generalizable to other species, we conducted a test where models were trained with human data and tested on mouse data unseen during model training, and vice versa (Figure 5A). The tests were performed with ChIP-seq and DNase-seq data, which was available for both species. Results showed that ChIP-seq models trained with a particular species can predict the quality of files from the same and the other unseen species with comparable performance. This could not be clearly observed with DNase-seq, for which prediction performance dropped substantially when trying to predict the file quality from data of unseen species. Therefore, cross-species generalizability of the models can be considered assay dependent. ROC-curves derived on the different feature sets are provided in supplementary (Figure S4).

**Figure 5.**
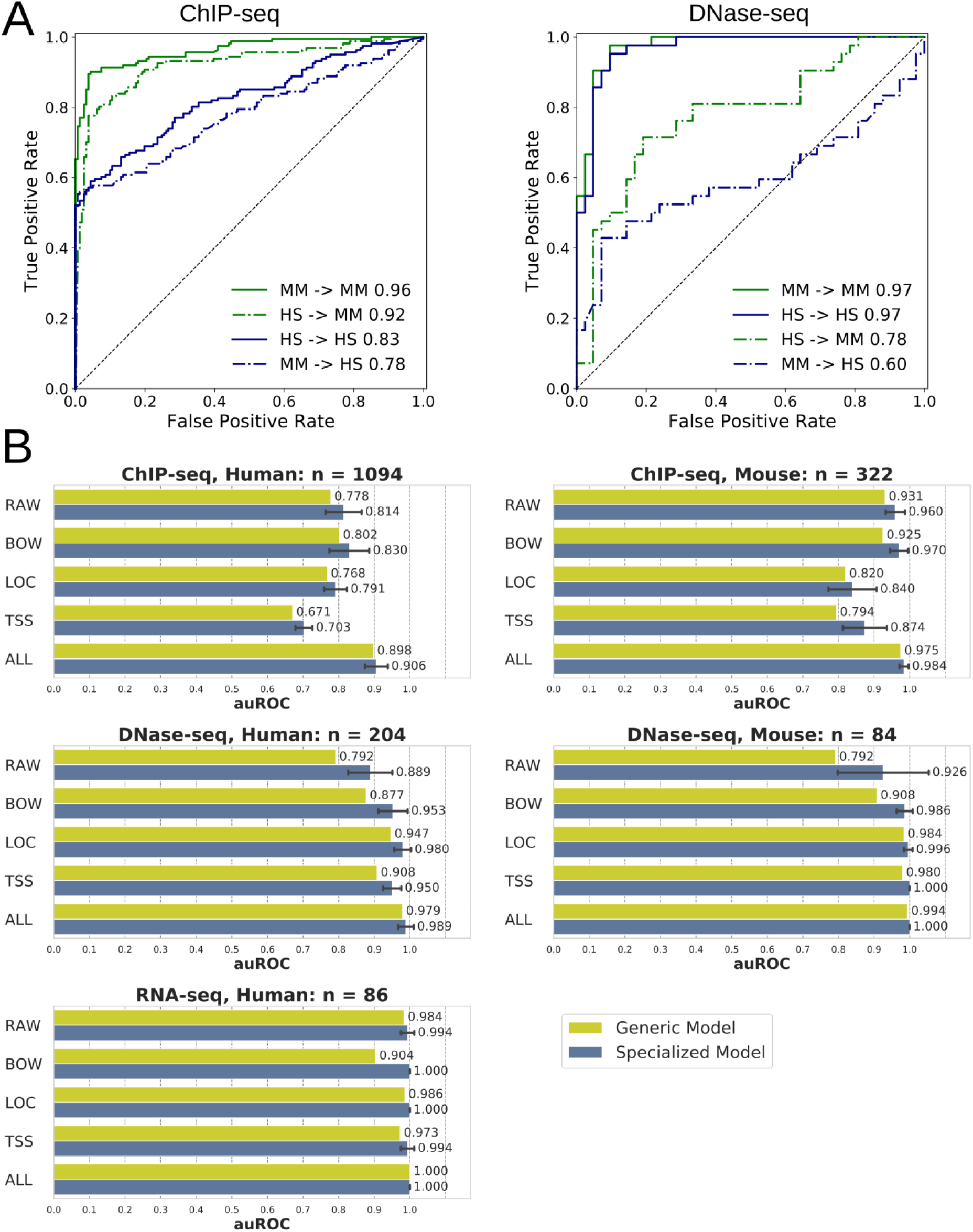
Unbiased and generalizable models. **A** Good performances of species-specific models in cross-species predictions demonstrate model generalization to other species. ROC curves show the classification performance for different species-assay combinations using all features. The solid lines represent cases in which data from the same species was used to define training and testing sets. The dashed lines show the performance on cases in which the species defining the training data differs from the species defining the testing data; e.g. “HS → MM” means the training set contains only data from human (Homo sapiens), while the testing data is only from mouse (Mus musculus). Legends also show corresponding auROC values. **B** Correlation of predictive performance of a generic model compared to different specialized models demonstrates lack of bias. For each feature set, performance of the generic model is detailed across the data subsets (green bars) and compared to specialized models trained for each combination of feature set and data subset (blue bars). Error bars show standard deviations derived from 10-fold cross-validations within the grid search. Feature sets: RAW (raw data), MAP (genome mapping), LOC (genomic localization), TSS (transcription start sites profile), ALL (all features). ROC: receiver operating characteristics. auROC: area under ROC curve.

### Generic model

A model that was of importance to us was the generic model trained on files from different species and different assays of the full dataset. Although its performance was high during the cross-validations (Figure 2) we were interested to know if it could be biased to data files from a particular subset, especially because the human ChIP-seq subset is overrepresented (52% of the dataset). After tuning and cross-validation of the generic model, we detailed its performance for each data subset (Figure 5B and S5B). The generic model was able to predict the quality of files from each subset with high performance. Still, specialized models, specifically trained on each subset, performed slightly better.

### Application to independent diagnostic studies

We have shown above that machine learning models trained on appropriate data from the same source are powerful and unbiased predictors of NGS file quality. Testing the models on independent datasets from another source such as the GEO database would allow their evaluation on different use cases including diagnosis and tell if more specific training would be required.

The potential of the models to filter low quality files from diagnostic studies was evaluated on two independent gene expression studies related to diseases. Assuming a large effect of the quality on gene expression data, being able to automatically identify files of low quality would potentially prevent patients from receiving inappropriate medications or treatments. The analysis of files from patients and control samples from the studies showed that the model outcome was not randomly associated with the samples. In the hepatocellular carcinoma and fatty liver studies, samples predicted to be of the lowest quality were farther from the cluster center of their group on a two-dimensional projection (Figure 6A and 6B). These samples are potentially outliers that should be reviewed to decide their relevance for downstream analysis. As we have observed that probabilities returned by the models may not be directly comparable between models and studies (data not shown), we would recommend for similar application to train the models on clinical data labelled as low- and high-quality when available.

**Figure 6.**
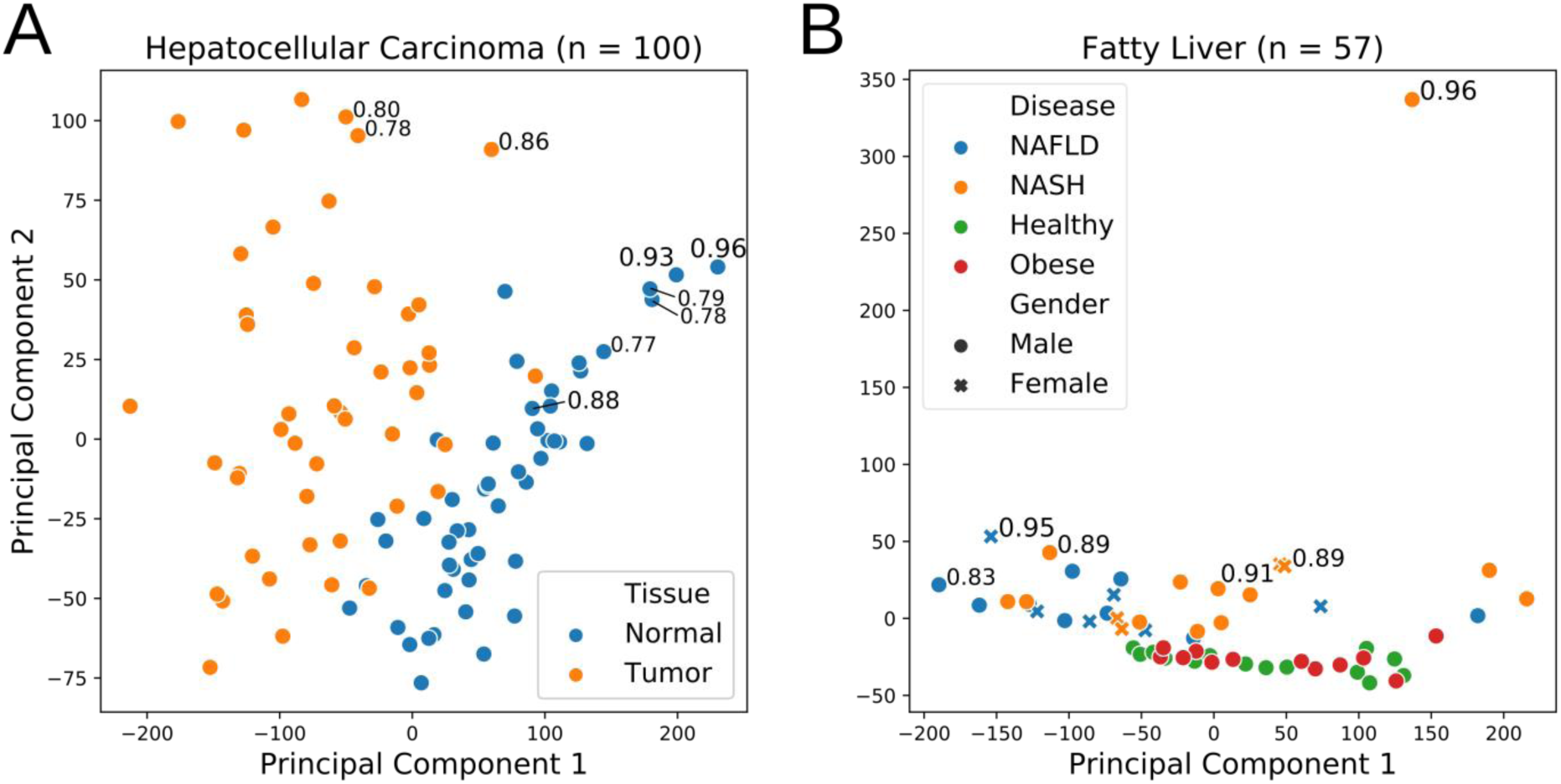
Identifying outliers in diagnostic studies. Gene expression data of human disease samples (RNA-seq) were retrieved from two independent diagnosis studies in the GEO database: (**A**) hepatocellular carcinoma and (**B**) fatty liver. For each study, samples were plotted using the 2 first components of a principal component analysis applied on gene expression profiles. For 10% of samples with the highest P_low_ (predictive probability to be of low-quality) the probability is shown next to the sample dot.

## Discussion

The versatility and power of NGS applications makes the sequencing technology a popular tool in biology and medicine. The complexity to evaluate data quality leads to non-optimal data file filtering and consequently has a negative impact on research and clinical results. Using thousands of files from ENCODE, we first derived statistics-based guidelines to interpret NGS quality features from standard software tools. Then, using a systematic method testing 10 algorithms, different sets of features and numerous parameters, we were able to build unbiased optimal models able to accurately predict the quality of NGS data files. The classification models outperformed the baseline (the best single feature) and were applicable to other assays and species. Application of this method on labelled database content and disease datasets was very promising in its ability to clearly identify problematic samples.

The study of quality features derived by widely used bioinformatic tools on the ENCODE files provides statistic-based guidelines to NGS specialists who have to make a manual decision of high complexity on the quality of their files (Figure S1). For instance, FastQC is the most widely used tool to decide the quality of all types of NGS assays. Yet, taken independently, its derived quality features show poor or moderate performance in predicting RNA-seq or DNAse-seq file quality, some of the features being not informative in any situation. On the contrary, other features and tools are more recommendable and, thanks to our guidelines, NGS specialists will know exactly if their files are more comparable to high- or low-quality files from ENCODE.

The best individual features were systematically outperformed by machine learning models that combine multiple features. Such algorithms, especially from the field of deep learning, benefit from bigger datasets but we were not able to find a data repository comparable to our main source (ENCODE) for integration. The restricted amount of low-quality data in ENCODE was therefore the main limitation of our study that could have generated biases and non-generalizable models. Our models have been trained only on two species and three assays for which we found a reasonable number of files. Interestingly, the generalization across different species of some models overcome this limitation and thanks to our systematic evaluations we found no biases towards data subsets or experiments. Nevertheless, specialized models trained on data subsets, such as human ChIP-seq or mouse DNAse-seq, performed slightly better than a generic model. Models may also benefit from additional features from other NGS software tools or genomic annotations [31], [32]. As it has been noted in other contexts, negative results can be valuable [33], and we encourage researchers that generate new data to share negative results or low-quality NGS files with the community together with high-quality files in order to enable more accurate and more generalizable models for NGS quality control.

The possible applications detailed in this study highlight the usefulness of our predictive models. Either as a researcher or as a database curator who wishes to identify low-quality files, using the models as decision support tool can save a substantial amount of resources (up to 50% for problematic experiments). For selected external disease datasets, models trained with ENCODE data have shown their relevance by classifying true and potential outlier samples. Nevertheless, as explained above, we would highly recommend to directly train models on more similar clinical data to create the best possible models that will prevent the impact of quality issues to diagnosis and therefore to enable patients to receive appropriate medication or treatment. Calibration methods may also be used to better compare results from different models [28]. Finally, results obtained on samples undergoing major DNA damage or rearrangements, induced by particular cancer cells for example, should be taken carefully as models would be limited to features depending on a healthy reference genome and may confuse these phenomena with quality issues.

## Conclusions

We have statistically characterized common NGS quality features of a large set of data files and optimized their complex quality control using a machine learning approach. The derived statistical guidelines and predictive models represent a valuable resource for NGS specialists. Predictive models are unbiased, accurate and to some extent widely applicable to unseen data types. Given enough labelled data for training, this approach could work for any type of NGS assay or species. Therefore, we strongly encourage researchers to share both high- and low-quality data with the community.

## Methods

### Dataset

To analyze the potential of machine learning applied to quality assessment of NGS data files, we implemented a workflow as shown in (Figure 1A). FastQ files and quality annotations were downloaded from the ENCODE data portal. The ENCODE status represents the result of a comprehensive manual inspection of the data by scientists from the Encode’s Data Coordination Center according to the ENCODE guidelines with a main focus on the quality. FastQ files that are uploaded to ENCODE and assigned to an experiment are initially released. When serious issues, mostly because of insufficient quality, are recognized, whole experiment or single files are revoked. We use this status as a good indication for the quality of a file and considered FastQ files as of low-quality when revoked and high-quality when released.

We downloaded 1321 low-quality files plus the same number of high-quality files (total = 2642) to define balanced training and testing sets. For the selection of high-quality files, we prioritized files that are associated with an experiment that contains also revoked files. The remaining high-quality files were chosen randomly. From this full dataset, we defined data subsets as sets of files representing a combination of species and assay such as human ChIP-seq or mouse DNAse-seq (Figure 1B). Because of their higher number of files and to facilitate comparisons between species, we used the following subsets for training machine learning models: human ChIP-seq (single-end and paired-end), mouse ChIP-seq (single-end), human and mouse DNAse-seq (paired-end), and human RNA-seq. ChIP-seq results are discussed in the article and plotted in figures only for single-end files (a dedicated supplementary figure shows results for paired-end files).

### Deriving Quality Feature Sets

We derived four different feature sets for the set of downloaded FastQ files as visualized in the sub-workflow in (Figure 1A). The first feature set RAW was defined by eleven features from the summary statistics of the FastQC tool (http://www.bioinformatics.babraham.ac.uk/projects/fastqc/). In the summary of a FastQC report, each statistic is flagged as Fail, Warning, or Pass. We use these flags as values for the features. The second feature set MAP contained the mapping statistics after applying Bowtie2 [5] to map the sequencing reads of human and mouse against the hg38 and mm10 genome assemblies, respectively. The mapping statistics describe the percentage of reads being unmapped, uniquely mapped, or multiply mapped and their overall mapping rate. Accordingly, there are four features for single-end and eight features for paired-end as these statistics are done for both the concordantly and discordantly mapped reads. The third feature set LOC is composed of nine features describing the distribution of reads mapped within the following types of genomic regions of interest: promoter, first intron, other introns, 5’UTR, first exon, other exons, 3’UTR, distal intergenic, and downstream proximal to the 3’UTR. The features were derived using the Bioconductor package ChIPseeker [22]. The fourth feature set TSS describes the distribution of reads near TSS (transcription start site) positions in the genome. The Bioconductor package ChIPpeakAnno [23] was used to compute the number of reads within the region 5kb up- and downstream the TSS divided into ten bins, resulting in ten features for TSS identified by their central coordinate (e.g. TSS −4500 denotes the genomic region with the following boundaries relative to TSSs: −5kb and −4kb). To reduce memory requirements during computation, the features for LOC and TSS were derived on one million mapped reads randomly sampled. For paired-end files, the RAW features were derived independently for each of the two files, while MAP, LOC, and TSS features were derived for the pair of files itself. In order to reduce redundancy in the dataset, we filtered out the RAW features for one member of each pair randomly. The largest files that were created within this data preprocessing are the FastQ and BAM files that sum up to a data set size of 5.6 TB and 2 TB, respectively.

### Machine learning models

We conducted classification experiments on either all samples of the dataset (generic models) or on data subsets containing samples for a particular combination of species and assay (specialized models) as defined above. Based on the different species-assays and feature set combinations, machine learning algorithms were applied to train models that classify the quality class defined from the ENCODE status. A comprehensive grid search was applied to find optimal algorithm and parameter setting for each subset. The algorithm set is listed within (Figure 4). In combination with parameter settings specific to each algorithm, a total of 19417 different models were trained and evaluated within the grid search for each classification case. Furthermore, for each parameter setting, we applied a feature selection method prior to the classification. The feature selection is based on chi-squared statistics and selects the top k features. We analyzed four values for k: 100% (no feature selection), 75%, 50%, or 25% of features in the given set of quality features.

The predictive performance was evaluated by the area under receiver operating characteristic curve (auROC). Within the grid search and feature selection, a ten-fold cross-validation was applied to evaluate the predictive performance. The entire grid search (including the preprocessing methods as well as the chi-squared statistics and nine of the classification algorithms) was implemented using the Python package scikit-learn [24], [25]. XGBoost algorithm was implemented using an external library [26].

### Cross-species generalization

In order to test the generalization of classification models to data from unseen species, we performed classification experiments using training and testing sets containing data from different species, respectively. We used human and mouse ChIP-seq and DNase-seq data. Using a five-fold cross-validation, five training and testing sets were created for each species and the grid search was applied to the training sets. The best performing model for each training set was identified by the highest auROC achieved within a ten-fold cross-validation on the training set. Finally, each of these models was evaluated twice, firstly on the corresponding testing set from the five-fold cross-validation that contains data from the same species and secondly on the corresponding testing set containing data from the differing species.

### Human diagnostic studies

For this analysis we used three published datasets downloaded from GEO (Gene Expression Omnibus) [27], [28]. The data was preprocessed based on the workflow that we also applied on the data from ENCODE as explained above and in Figure 1A (extraction of quality feature sets). The mapped reads that were created within the workflow were used to quantify the gene expression counts using the function featureCounts from the Bioconductor package Rsubread [29]. The counts were normalized using TPM [30]. Finally, for all the three GEO datasets, gene expression values were log2 transformed and standardized before applying the Principal Component Analysis.

## References

[1] G. A. Merino, C. Fresno, F. Netto, E. D. Netto, L. Pratto, and E. A. Fernández, “The impact of quality control in RNA-seq experiments,” in Journal of Physics: Conference Series, 2016, vol. 705, no. 1, p. 12003.

[2] C. R. Williams, A. Baccarella, J. Z. Parrish, and C. C. Kim, “Trimming of sequence reads alters RNA-Seq gene expression estimates,” BMC Bioinformatics, vol. 17, no. 1, p. 103, 2016.

[3] S.-F. Yang, C.-W. Lu, C.-T. Yao, and C.-M. Hung, “To Trim or Not to Trim: Effects of Read Trimming on the De Novo Genome Assembly of a Widespread East Asian Passerine, the Rufous-Capped Babbler (Cyanoderma ruficeps Blyth),” Genes (Basel)., vol. 10, p. 737, 2019.

[4] C. A. Meyer and X. S. Liu, “Identifying and mitigating bias in next-generation sequencing methods for chromatin biology,” Nat. Rev. Genet., vol. 15, no. 11, pp. 709–721, 2014.

[5] B. Langmead and S. L. Salzberg, “Fast gapped-read alignment with Bowtie 2,” Nat. Methods, vol. 9, no. 4, p. 357, 2012.

[6] H. Li and R. Durbin, “Fast and accurate short read alignment with Burrows–Wheeler transform,” Bioinformatics, vol. 25, no. 14, pp. 1754–1760, 2009.

[7] A. Dobin et al., “STAR: ultrafast universal RNA-seq aligner,” Bioinformatics, vol. 29, no. 1, pp. 15–21, 2012.

[8] C. Trapnell, L. Pachter, and S. L. Salzberg, “TopHat: discovering splice junctions with RNA-Seq,” Bioinformatics, vol. 25, no. 9, pp. 1105–1111, 2009.

[9] M. D. Chikina and O. G. Troyanskaya, “An effective statistical evaluation of ChIPseq dataset similarity,” Bioinformatics, vol. 28, no. 5, pp. 607–613, 2012.

[10] M. I. Love, W. Huber, and S. Anders, “Moderated estimation of fold change and dispersion for RNA-seq data with DESeq2,” Genome Biol., vol. 15, no. 12, p. 550, 2014.

[11] G. K. Marinov, A. Kundaje, P. J. Park, and B. J. Wold, “Large-scale quality analysis of published ChIP-seq data,” G3 Genes, Genomes, Genet., vol. 4, no. 2, pp. 209–223, 2014.

[12] M.-A. Mendoza-Parra, W. Van Gool, M. A. Mohamed Saleem, D. G. Ceschin, and H. Gronemeyer, “A quality control system for profiles obtained by ChIP sequencing,” Nucleic Acids Res., vol. 41, no. 21, pp. e196.-e196, 2013.

[13] J. Brown, M. Pirrung, and L. A. McCue, “FQC Dashboard: integrates FastQC results into a web-based, interactive, and extensible FASTQ quality control tool,” Bioinformatics, vol. 33, no. 19, pp. 3137–3139, 2017.

[14] P. Ewels, M. Magnusson, S. Lundin, and M. Käller, “MultiQC: summarize analysis results for multiple tools and samples in a single report,” Bioinformatics, vol. 32, no. 19, pp. 3047–3048, 2016.

[15] A. Brazma, “Minimum information about a microarray experiment (MIAME)--successes, failures, challenges,” Sci. World J., vol. 9, pp. 420–423, 2009.

[16] J. Malone et al., “Modeling sample variables with an Experimental Factor Ontology,” Bioinformatics, vol. 26, no. 8, pp. 1112–1118, 2010.

[17] C. A. Davis et al., “The Encyclopedia of DNA elements (ENCODE): data portal update,” Nucleic Acids Res., vol. 46, no. D1, pp. D794–D801, 2017.

[18] “The ENCODE (ENCyclopedia Of DNA Elements) Project,” Science (80-.)., vol. 306, no. 5696, pp. 636–640, 2004.

[19] E. P. Consortium and others, “A user’s guide to the encyclopedia of DNA elements (ENCODE),” PLoS Biol., vol. 9, no. 4, p. e1001046, 2011.

[20] S. G. Landt et al., “ChIP-seq guidelines and practices of the ENCODE and modENCODE consortia,” Genome Res., vol. 22, no. 9, pp. 1813–1831, 2012.

[21] J. Li et al., “ForestQC: quality control on genetic variants from next-generation sequencing data using random forest,” bioRxiv, p. 444828, 2018.

[22] G. Yu, L.-G. Wang, and Q.-Y. He, “ChIPseeker: an R/Bioconductor package for ChIP peak annotation, comparison and visualization,” BMC Bioinformatics, vol. 31, no. 14, pp. 2382–2383, 2015.

[23] L. J. Zhu et al., “ChIPpeakAnno: a Bioconductor package to annotate ChIP-seq and ChIP-chip data,” BMC Bioinformatics, vol. 11, no. 1, p. 237, 2010.

[24] L. Buitinck et al., “{API} design for machine learning software: experiences from the scikit-learn project,” in ECML PKDD Workshop: Languages for Data Mining and Machine Learning, 2013, pp. 108–122.

[25] F. Pedregosa et al., “Scikit-learn: Machine Learning in {P}ython,” J. Mach. Learn. Res., vol. 12, pp. 2825–2830, 2011.

[26] T. Chen and C. Guestrin, “{XGBoost}: A Scalable Tree Boosting System,” in Proceedings of the 22nd ACM SIGKDD International Conference on Knowledge Discovery and Data Mining, 2016, pp. 785–794.

[27] G. Liu, G. Hou, L. Li, Y. Li, W. Zhou, and L. Liu, “Potential diagnostic and prognostic marker dimethylglycine dehydrogenase (DMGDH) suppresses hepatocellular carcinoma metastasis in vitro and in vivo,” Oncotarget, vol. 7, no. 22, p. 32607, 2016.

[28] M. P. Suppli et al., “Hepatic transcriptome signatures in patients with varying degrees of nonalcoholic fatty liver disease compared with healthy normal-weight individuals,” Am. J. Physiol. Liver Physiol., vol. 316, no. 4, pp. G462--G472, 2019.

[29] Y. Liao, G. K. Smyth, and W. Shi, “The R package Rsubread is easier, faster, cheaper and better for alignment and quantification of RNA sequencing reads,” Nucleic Acids Res., vol. 47, no. 8, pp. e47–e47, 2019.

[30] G. P. Wagner, K. Kin, and V. J. Lynch, “Measurement of mRNA abundance using RNA-seq data: RPKM measure is inconsistent among samples,” Theory Biosci., vol. 131, no. 4, pp. 281–285, 2012.

[31] S. W. Wingett and S. Andrews, “FastQ Screen: A tool for multi-genome mapping and quality control,” F1000Research, vol. 7, 2018.

[32] H. M. Amemiya, A. Kundaje, and A. P. Boyle, “The ENCODE Blacklist: Identification of Problematic Regions of the Genome,” Sci. Rep., vol. 9, no. 1, p. 9354, 2019.

[33] P. Blohm et al., “Negatome 2.0: a database of non-interacting proteins derived by literature mining, manual annotation and protein structure analysis,” Nucleic Acids Res., vol. 42, no. D1, pp. D396–D400, 2013.

