## Supplementary Figures for "Automated quality control of next generation sequencing data using machine learning"

**Figure S1 – Statistical guidelines computed on the ENCODE files selection**

For each data subset, bar plots or boxplots show the distributions of (**A**) the raw data features derived by the FastQC tool, (**B**) the genome mapping features (MAP) derived by Bowtie2, (**C)** the genomic localization features (LOC) derived by the ChIPSeeker R library, (**D)** the transcription start sites profile features (TSS) derived by the ChIPpeakAnno R library.

A

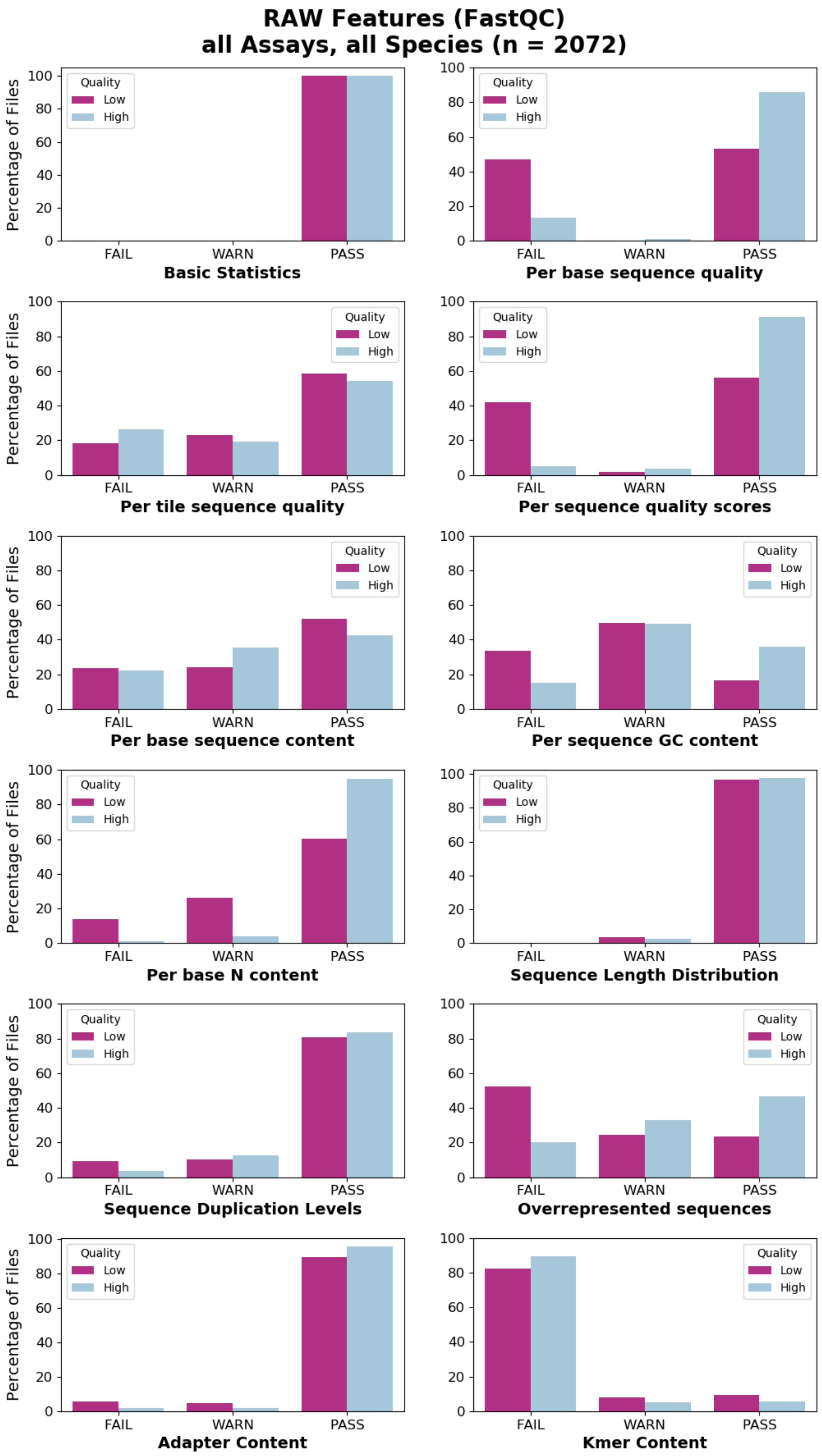
RAW Features (FastQC)
all Assays, all Species (n = 2072)
Percentage of Files
Quality
Low
High
FAIL
WARN
PASS
Basic Statistics
Per base sequence quality
Per tile sequence quality
Per sequence quality scores
Per base sequence content
Per sequence GC content
Per base N content
Sequence Length Distribution
Sequence Duplication Levels
Overrepresented sequences
Adapter Content
Kmer Content

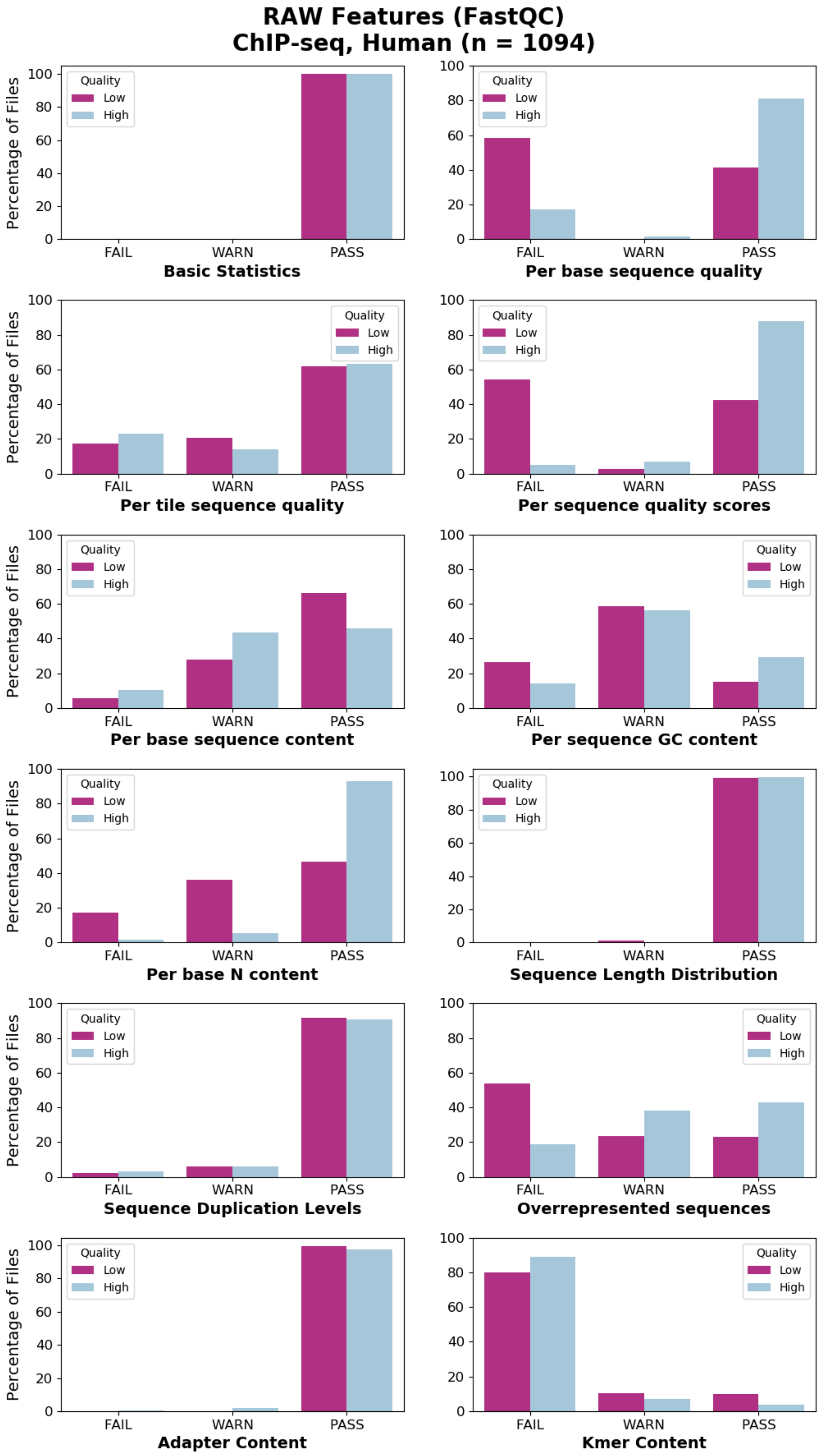
RAW Features (FastQC)
ChIP-seq, Human (n = 1094)
Percentage of Files
Quality
Low
High
FAIL
WARN
PASS
Basic Statistics
Per base sequence quality
Per tile sequence quality
Per sequence quality scores
Per base sequence content
Per sequence GC content
Per base N content
Sequence Length Distribution
Sequence Duplication Levels
Overrepresented sequences
Adapter Content
Kmer Content
0
20
40
60
80
100

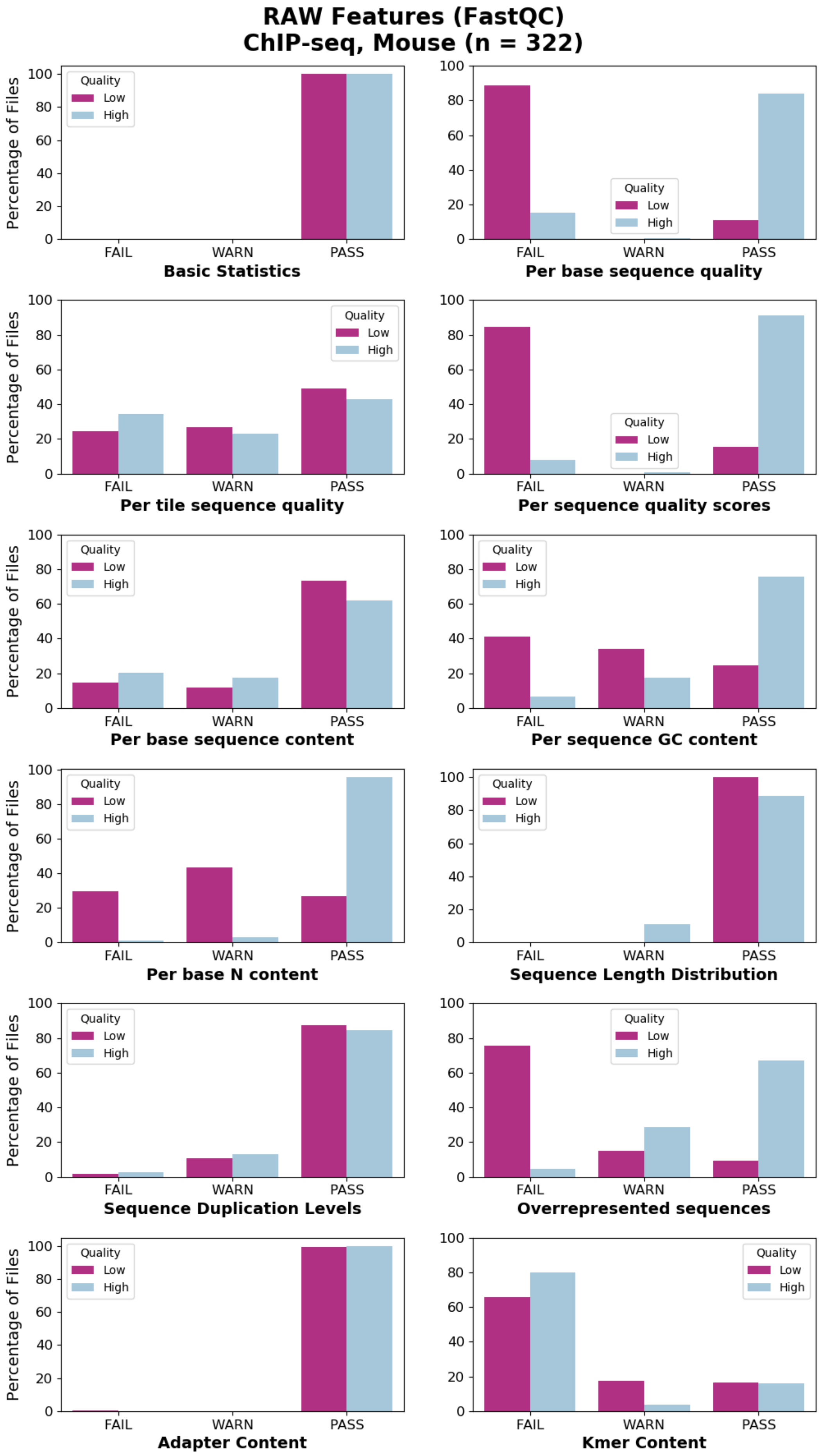
RAW Features (FastQC)
ChIP-seq, Mouse (n = 322)
Percentage of Files
Quality
Low
High
FAIL
WARN
PASS
0
20
40
60
80
100
Basic Statistics
Per base sequence quality
Per tile sequence quality
Per sequence quality scores
Per base sequence content
Per sequence GC content
Per base N content
Sequence Length Distribution
Sequence Duplication Levels
Overrepresented sequences
Adapter Content
Kmer Content

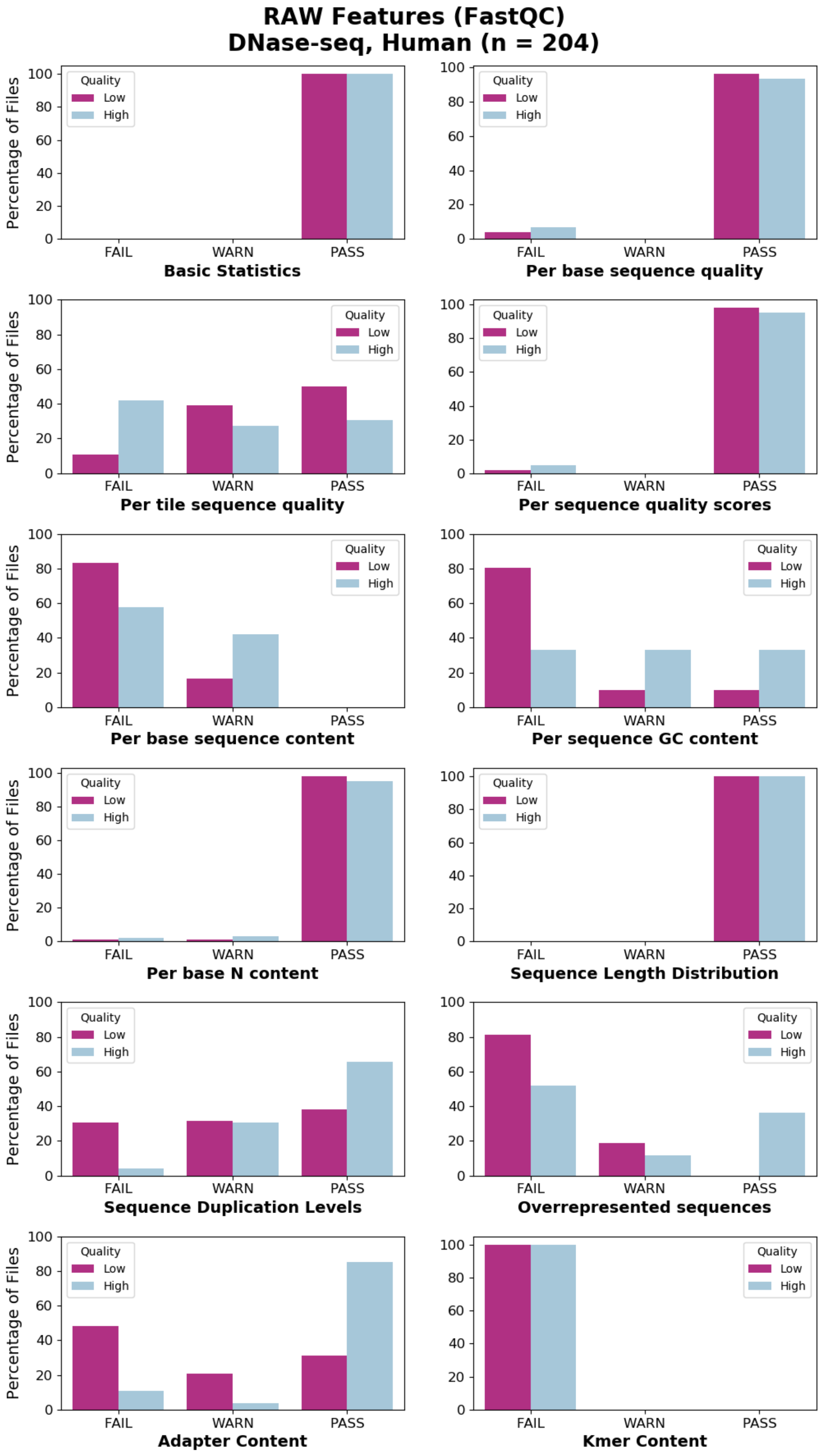
RAW Features (FastQC)
DNase-seq, Human (n = 204)
Percentage of Files
Quality
Low
High
FAIL
WARN
PASS
Basic Statistics
Per base sequence quality
Per tile sequence quality
Per sequence quality scores
Per base sequence content
Per sequence GC content
Per base N content
Sequence Length Distribution
Sequence Duplication Levels
Overrepresented sequences
Adapter Content
Kmer Content

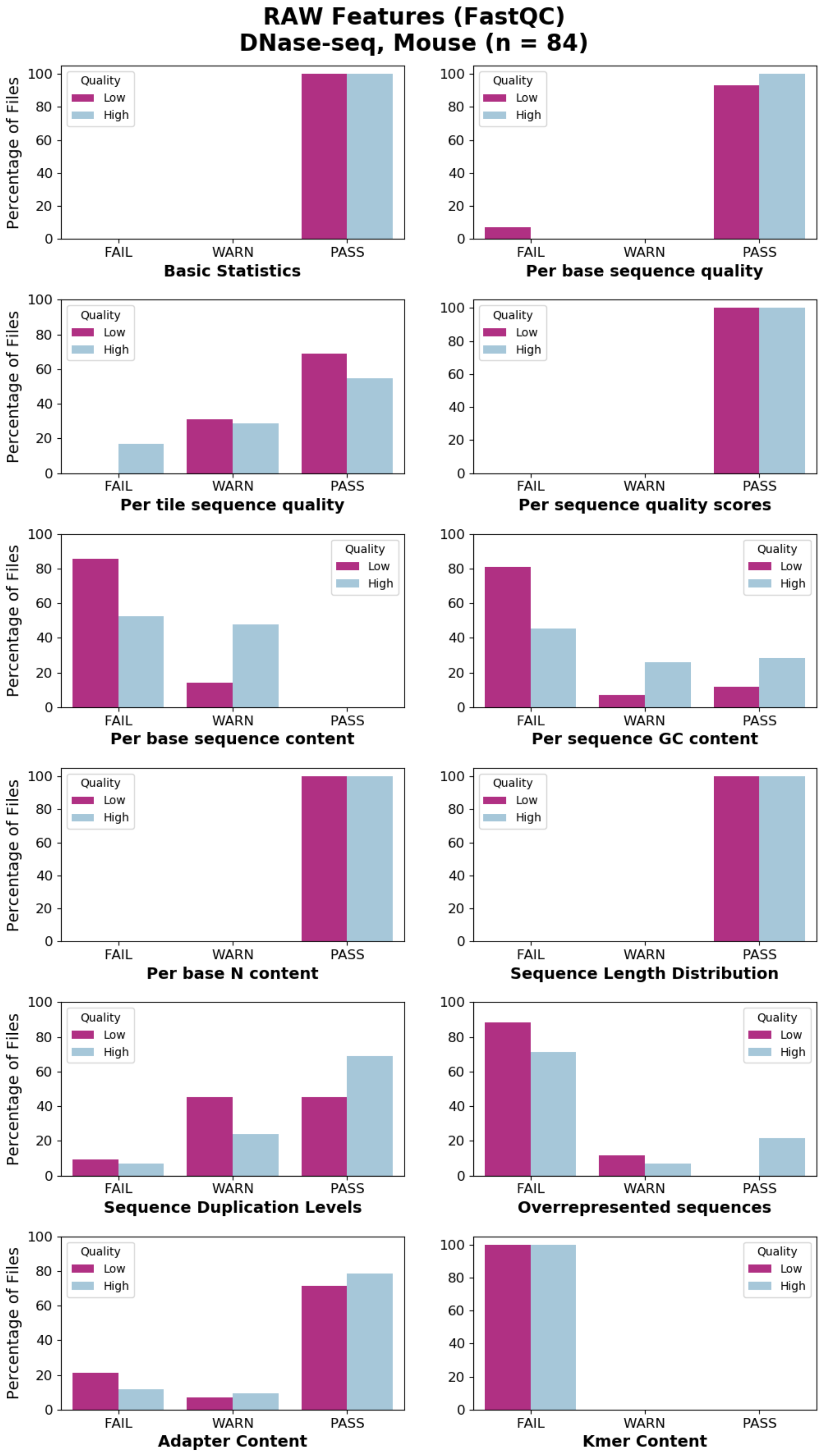
RAW Features (FastQC)
DNase-seq, Mouse (n = 84)
Percentage of Files
Quality
Low
High
FAIL
WARN
PASS
Basic Statistics
Per base sequence quality
Per tile sequence quality
Per sequence quality scores
Per base sequence content
Per sequence GC content
Per base N content
Sequence Length Distribution
Sequence Duplication Levels
Overrepresented sequences
Adapter Content
Kmer Content

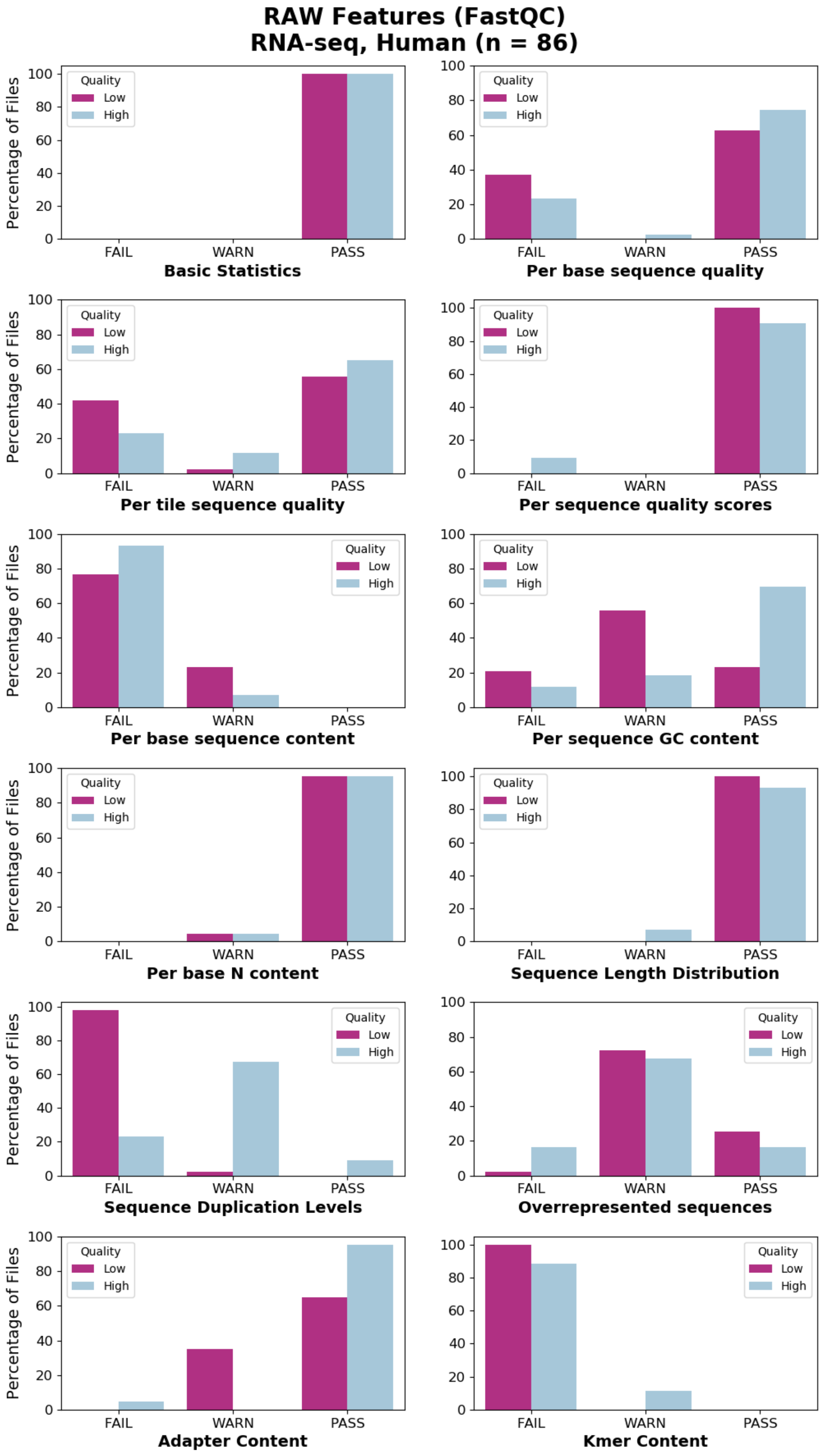
RAW Features (FastQC)
RNA-seq, Human (n = 86)
Percentage of Files
Quality
Low
High
FAIL
WARN
PASS
Basic Statistics
Per base sequence quality
Per tile sequence quality
Per sequence quality scores
Per base sequence content
Per sequence GC content
Per base N content
Sequence Length Distribution
Sequence Duplication Levels
Overrepresented sequences
Adapter Content
Kmer Content

B

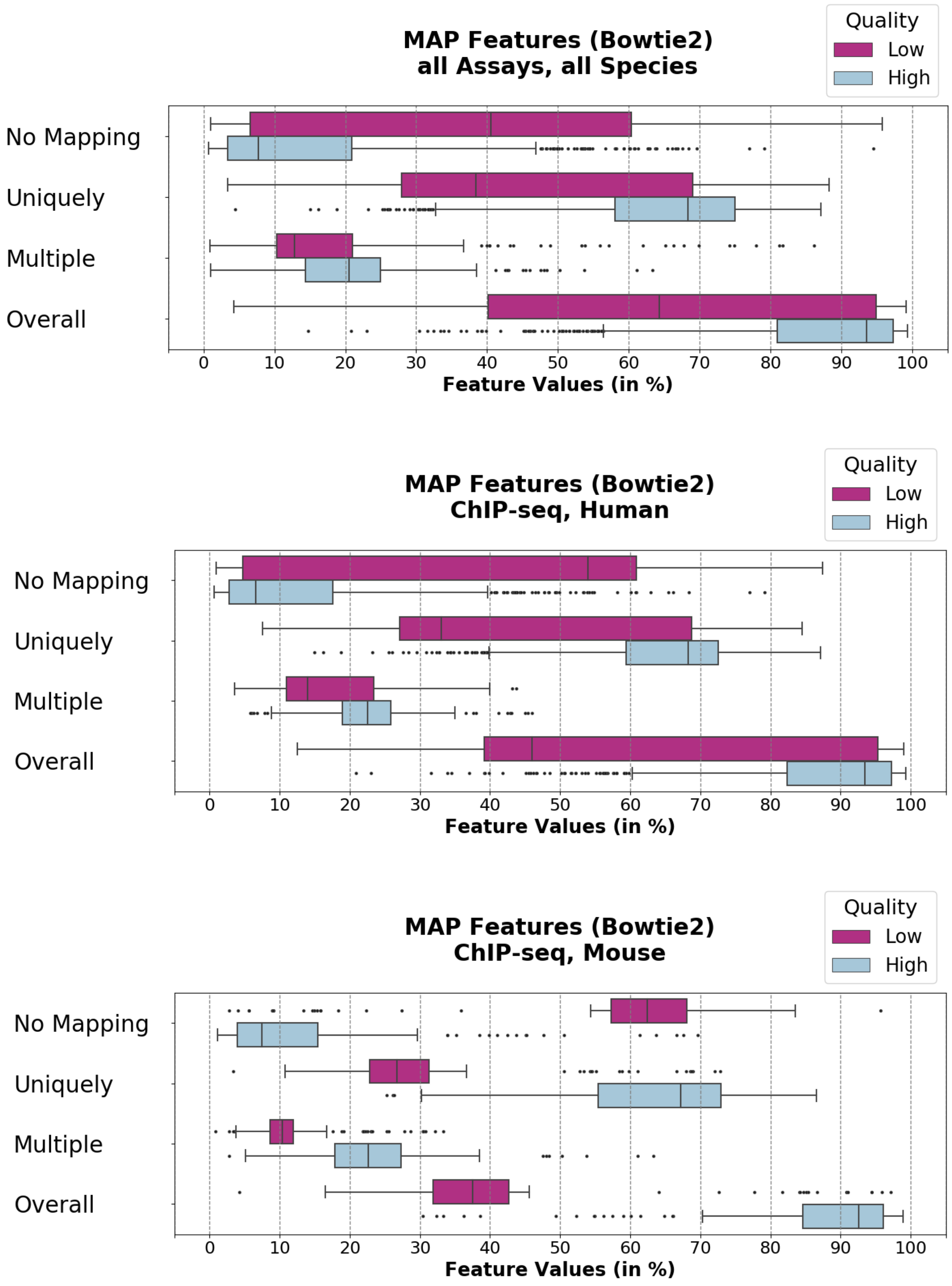

MAP Features (Bowtie2)
all Assays, all Species
Quality
Low
High
No Mapping
Uniquely
Multiple
Overall
0
10
20
30
40
50
60
70
80
90
100
Feature Values (in %)
MAP Features (Bowtie2)
ChIP-seq, Human
Quality
Low
High
No Mapping
Uniquely
Multiple
Overall
0
10
20
30
40
50
60
70
80
90
100
Feature Values (in %)
MAP Features (Bowtie2)
ChIP-seq, Mouse
Quality
Low
High
No Mapping
Uniquely
Multiple
Overall
0
10
20
30
40
50
60
70
80
90
100
Feature Values (in %)

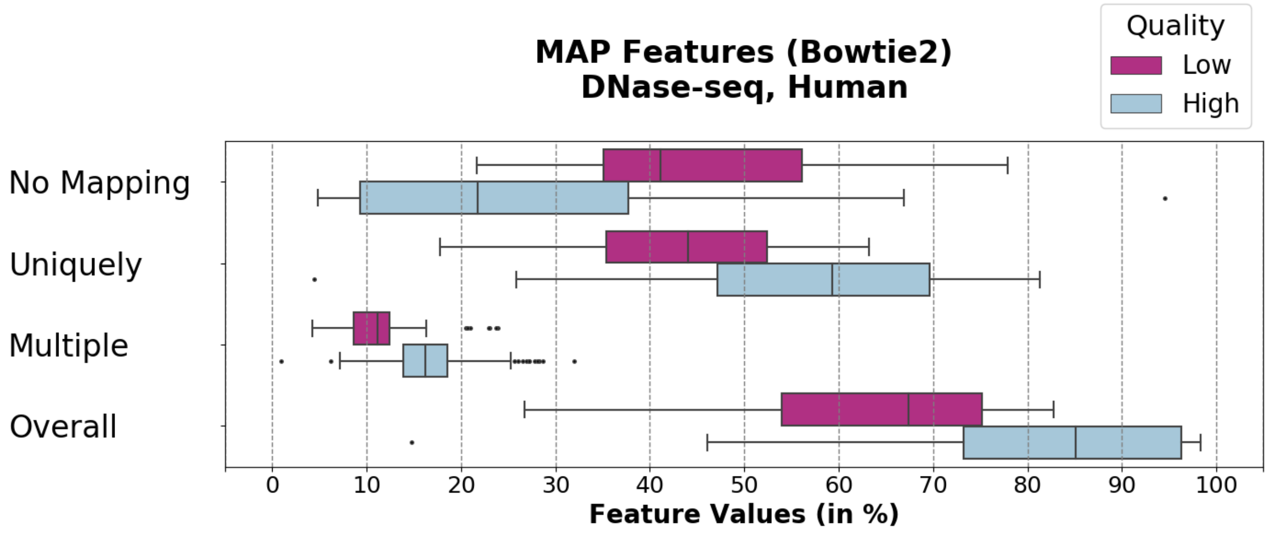
MAP Features (Bowtie2)
DNase-seq, Human
Quality
Low
High
No Mapping
Uniquely
Multiple
Overall
0
10
20
30
40
50
60
70
80
90
100
Feature Values (in %)

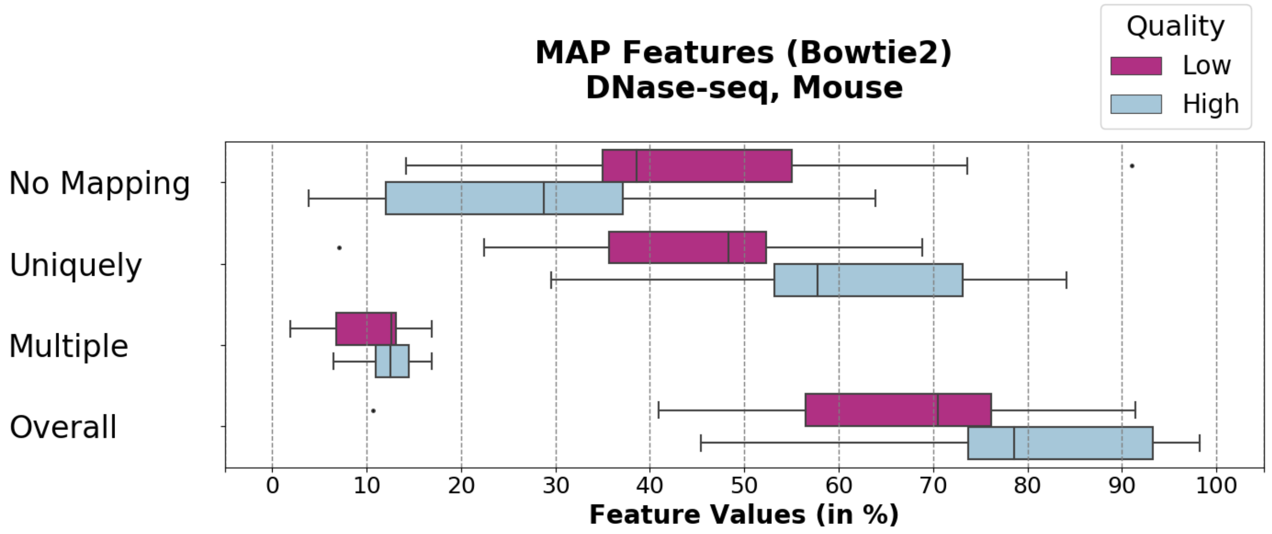
MAP Features (Bowtie2)
DNase-seq, Mouse
Quality
Low
High
No Mapping
Uniquely
Multiple
Overall
0
10
20
30
40
50
60
70
80
90
100
Feature Values (in %)

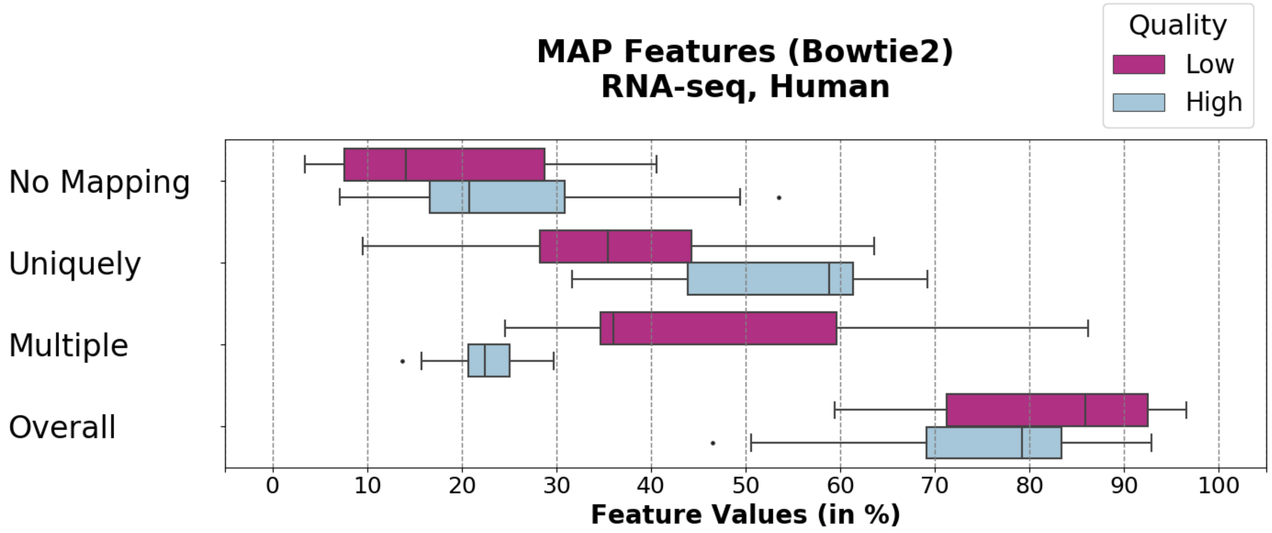
MAP Features (Bowtie2)
RNA-seq, Human
Quality
Low
High
No Mapping
Uniquely
Multiple
Overall
0
10
20
30
40
50
60
70
80
90
100
Feature Values (in %)

C

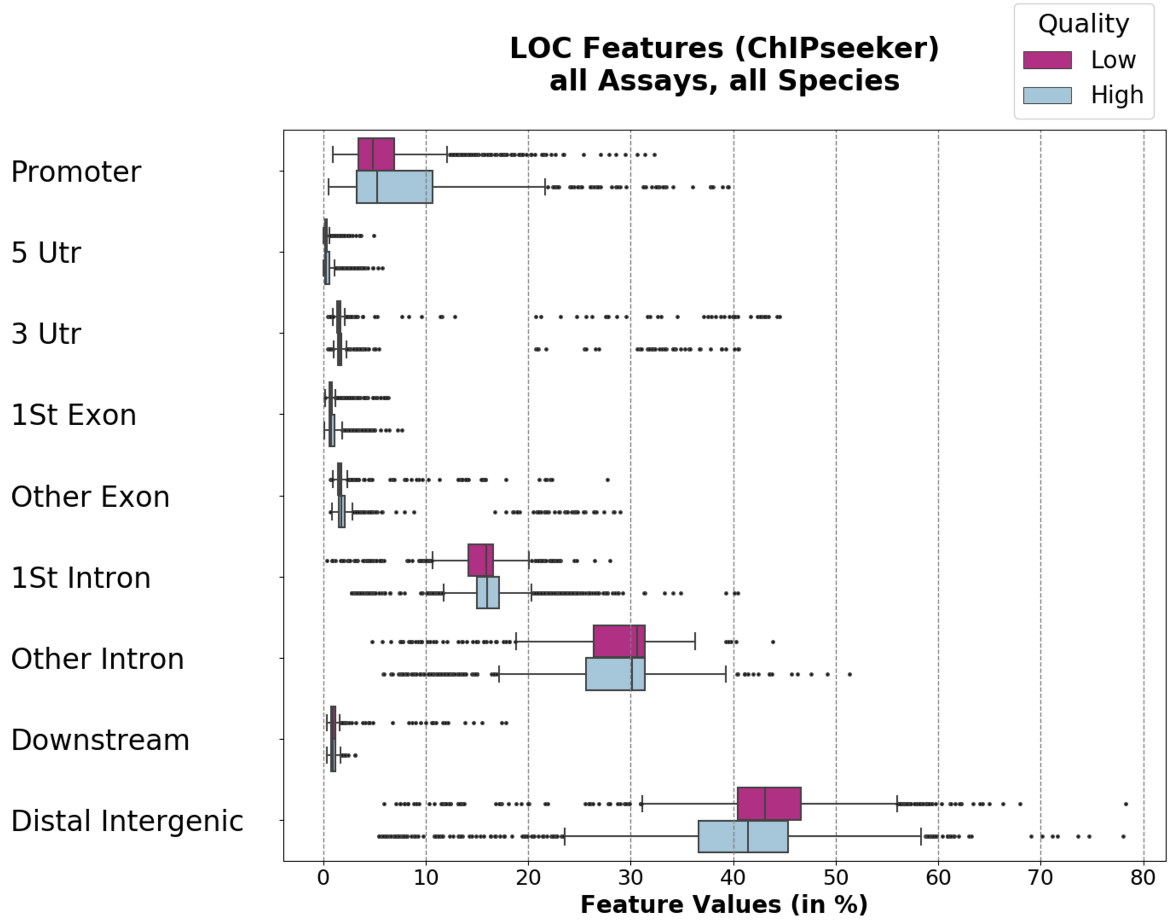

LOC Features (ChIPseeker)
all Assays, all Species
Quality
Low
High
Promoter
5 Utr
3 Utr
1St Exon
Other Exon
1St Intron
Other Intron
Downstream
Distal Intergenic
0
10
20
30
40
50
60
70
80
Feature Values (in %)


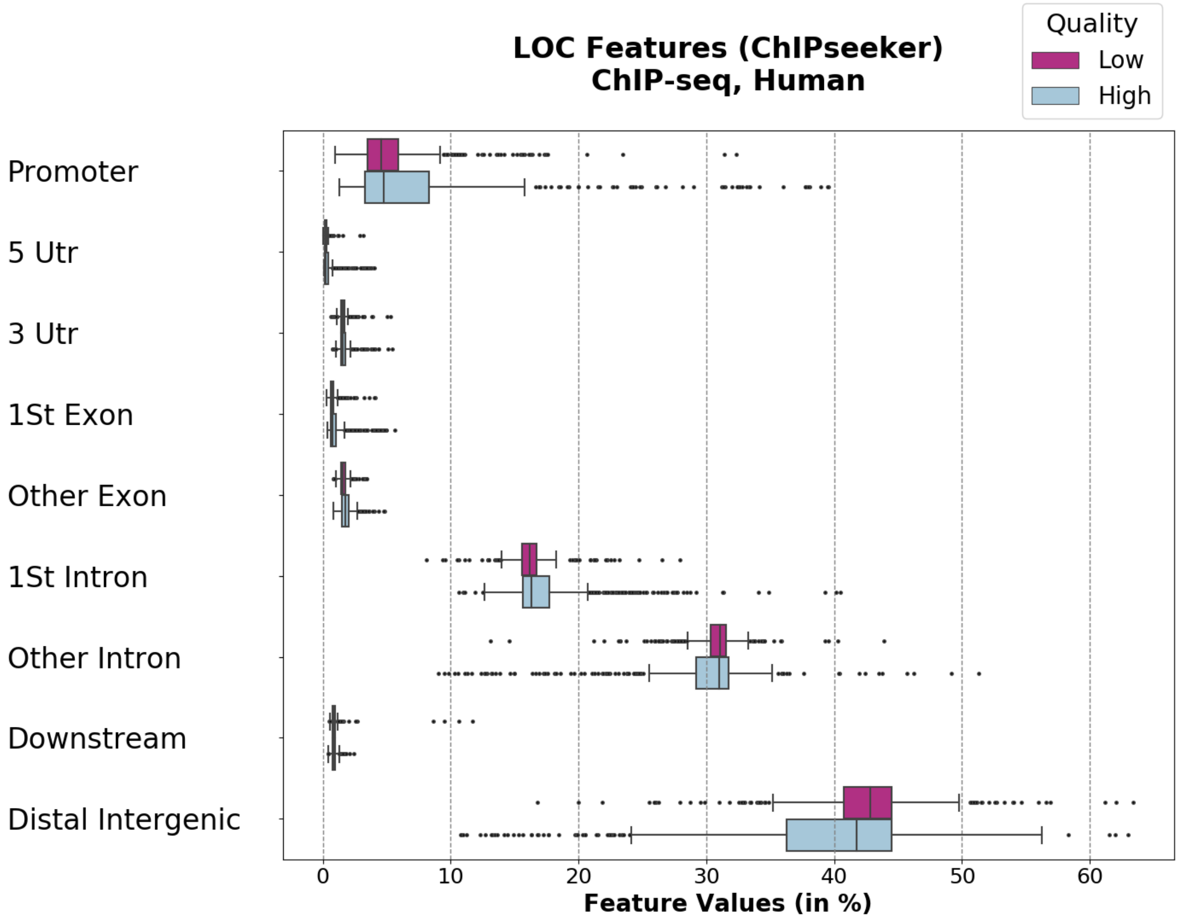

LOC Features (ChIPseeker)
ChIP-seq, Human
Quality
Low
High
Promoter
5 Utr
3 Utr
1St Exon
Other Exon
1St Intron
Other Intron
Downstream
Distal Intergenic
0
10
20
30
40
50
60
Feature Values (in %)

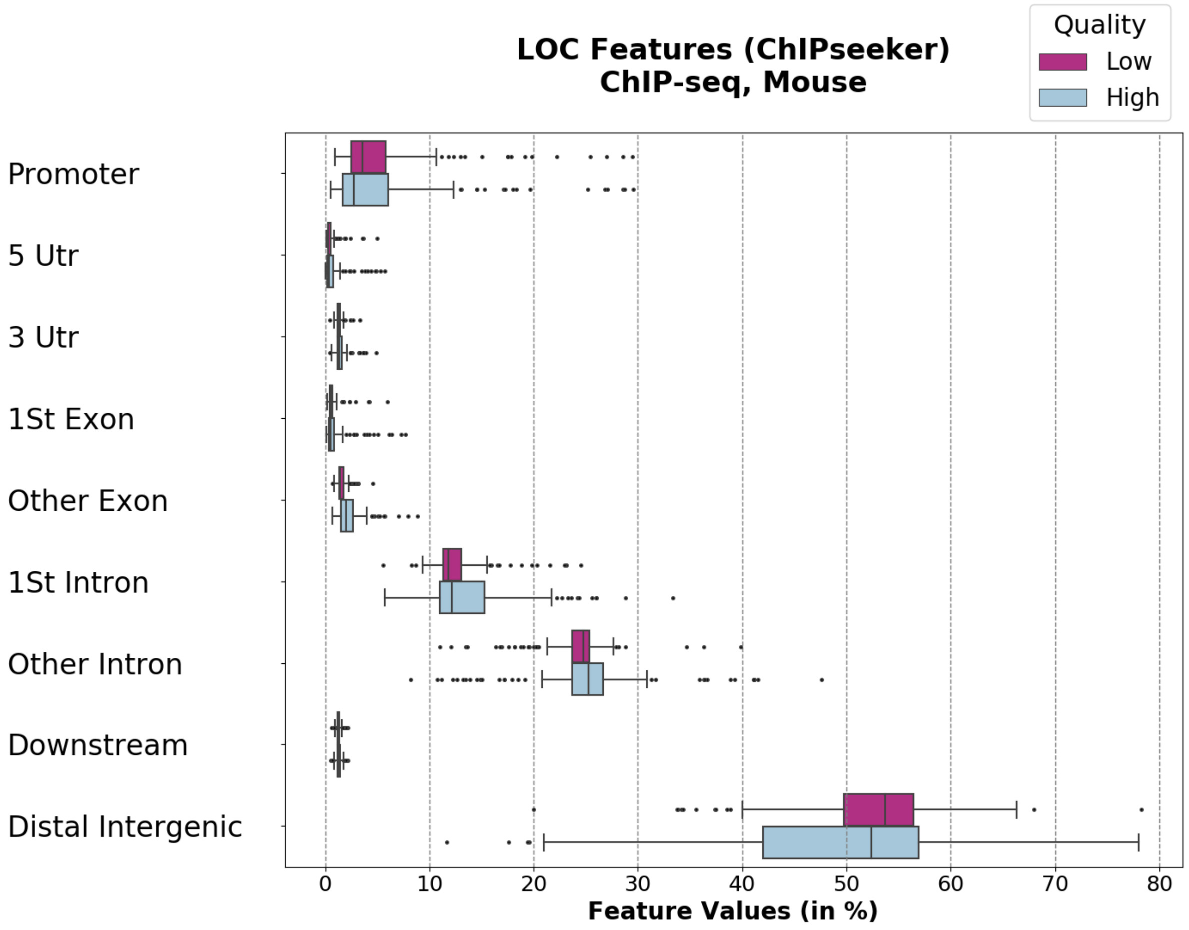

LOC Features (ChIPseeker)
ChIP-seq, Mouse
Quality
Low
High
Promoter
5 Utr
3 Utr
1St Exon
Other Exon
1St Intron
Other Intron
Downstream
Distal Intergenic
0
10
20
30
40
50
60
70
80
Feature Values (in %)


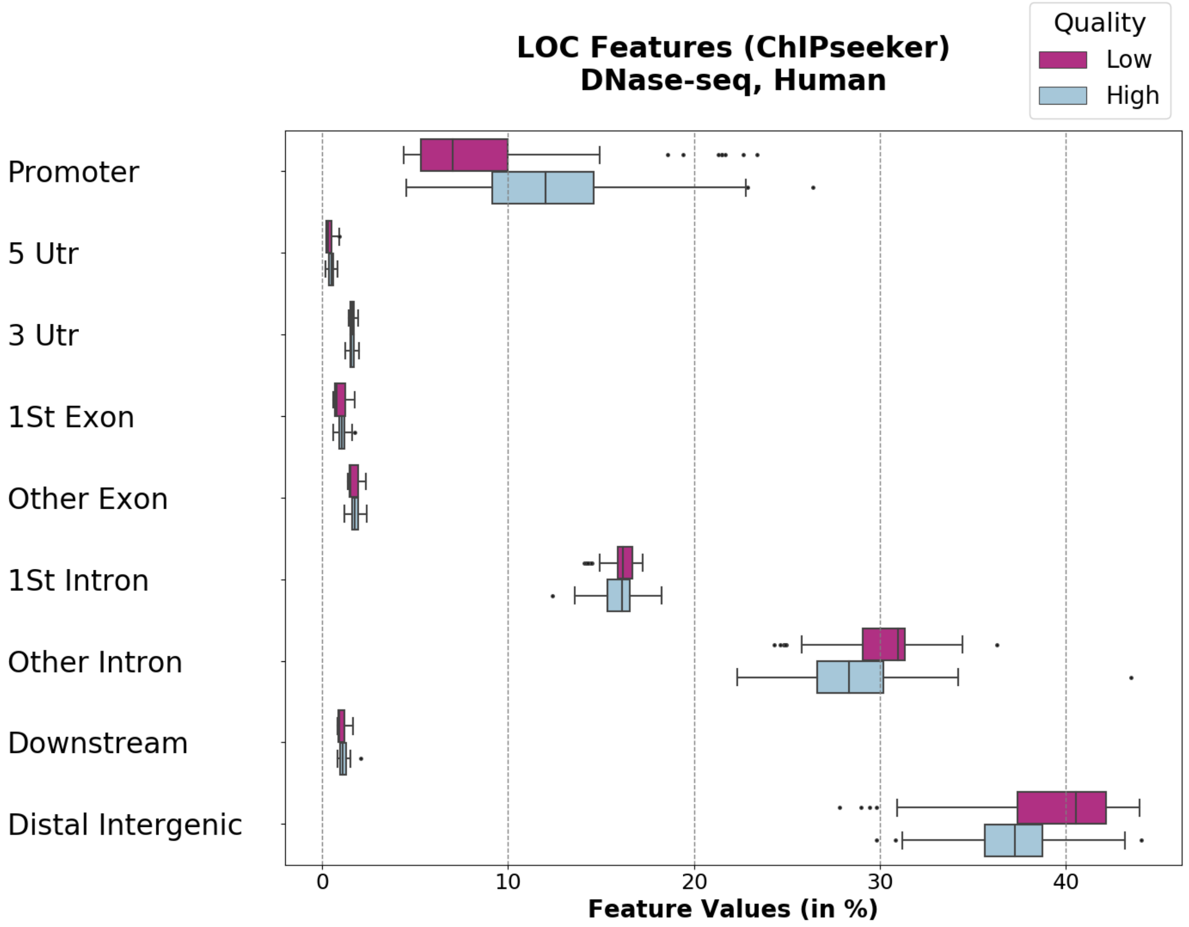

LOC Features (ChIPseeker)
DNase-seq, Human
Quality
Low
High
Promoter
5 Utr
3 Utr
1St Exon
Other Exon
1St Intron
Other Intron
Downstream
Distal Intergenic
0
10
20
30
40
Feature Values (in %)

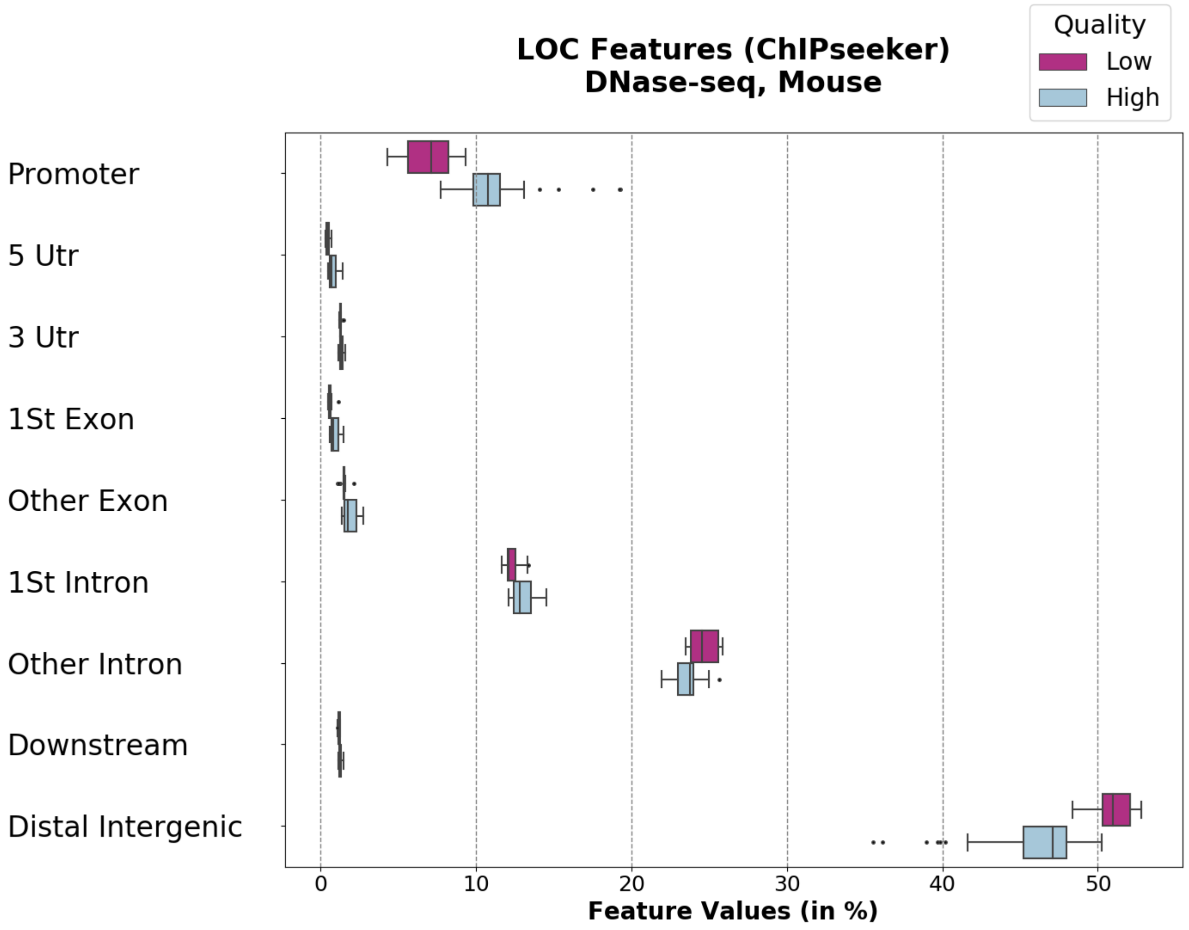
LOC Features (ChIPseeker)
DNase-seq, Mouse
Quality
Low
High
Promoter
5 Utr
3 Utr
1St Exon
Other Exon
1St Intron
Other Intron
Downstream
Distal Intergenic
0
10
20
30
40
50
Feature Values (in %)

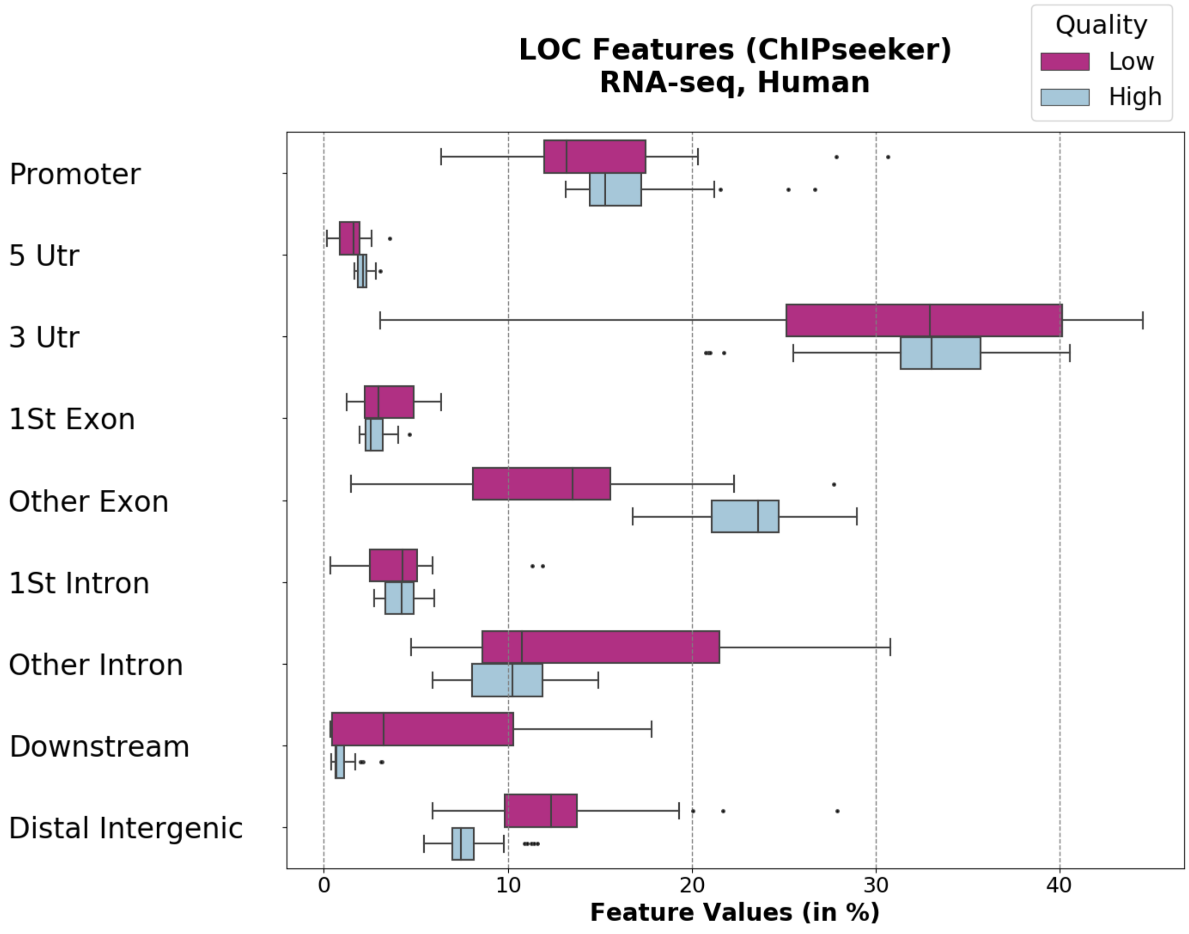
LOC Features (ChIPseeker)
RNA-seq, Human
Quality
Low
High
Promoter
5 Utr
3 Utr
1St Exon
Other Exon
1St Intron
Other Intron
Downstream
Distal Intergenic
0
10
20
30
40
Feature Values (in %)

**D**

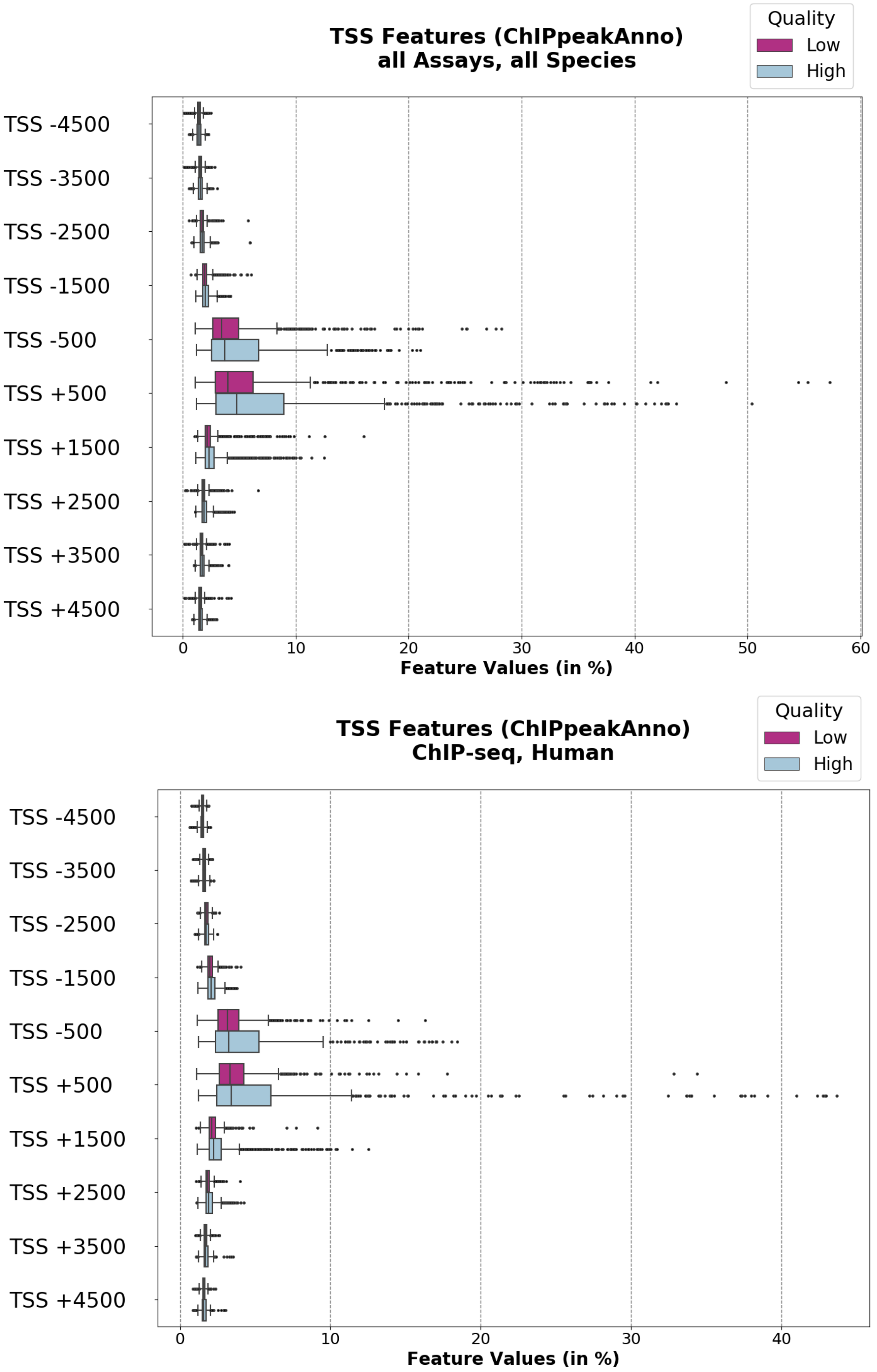
TSS Features (ChIPpeakAnno)
all Assays, all Species
Quality
Low
High
TSS -4500
TSS -3500
TSS -2500
TSS -1500
TSS -500
TSS +500
TSS +1500
TSS +2500
TSS +3500
TSS +4500
0
10
20
30
40
50
60
Feature Values (in %)
TSS Features (ChIPpeakAnno)
ChIP-seq, Human
Quality
Low
High
TSS -4500
TSS -3500
TSS -2500
TSS -1500
TSS -500
TSS +500
TSS +1500
TSS +2500
TSS +3500
TSS +4500
0
10
20
30
40
Feature Values (in %)

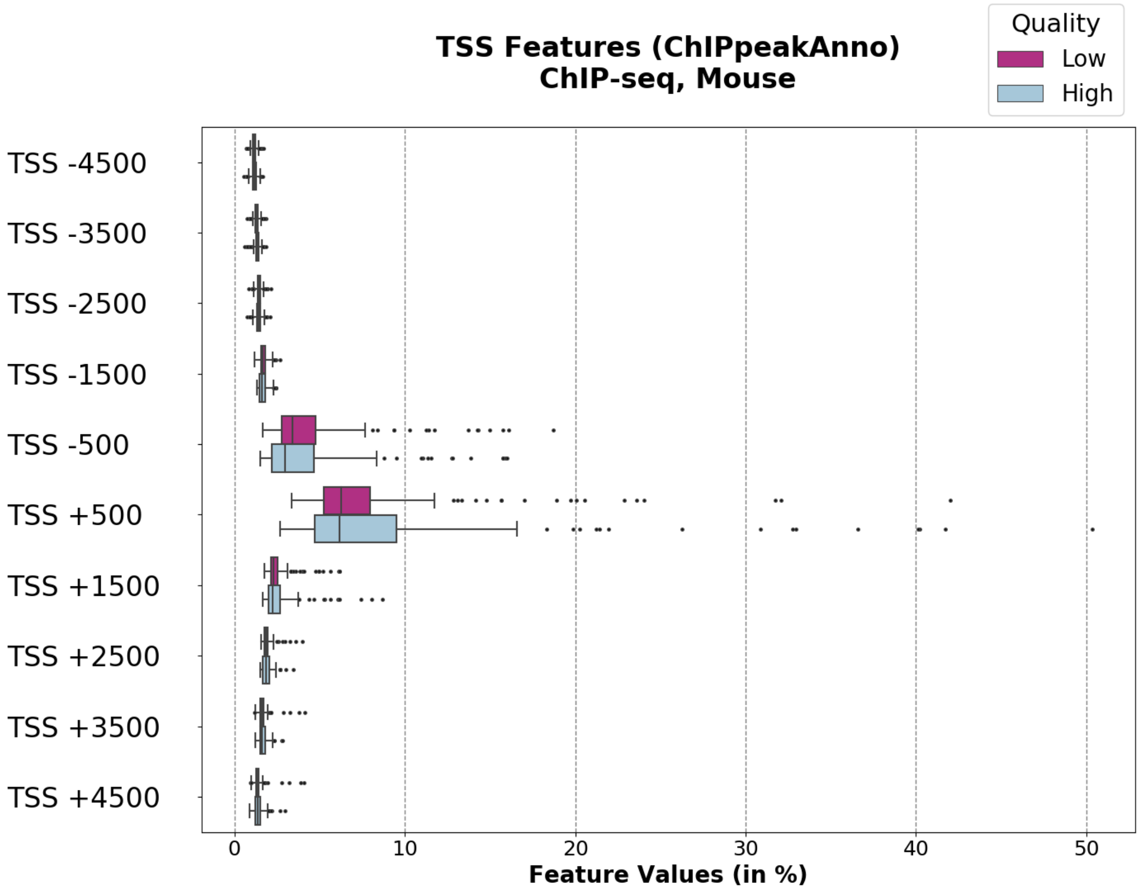
TSS Features (ChIPpeakAnno)
ChIP-seq, Mouse
Quality
Low
High
TSS -4500
TSS -3500
TSS -2500
TSS -1500
TSS -500
TSS +500
TSS +1500
TSS +2500
TSS +3500
TSS +4500
0
10
20
30
40
50
Feature Values (in %)

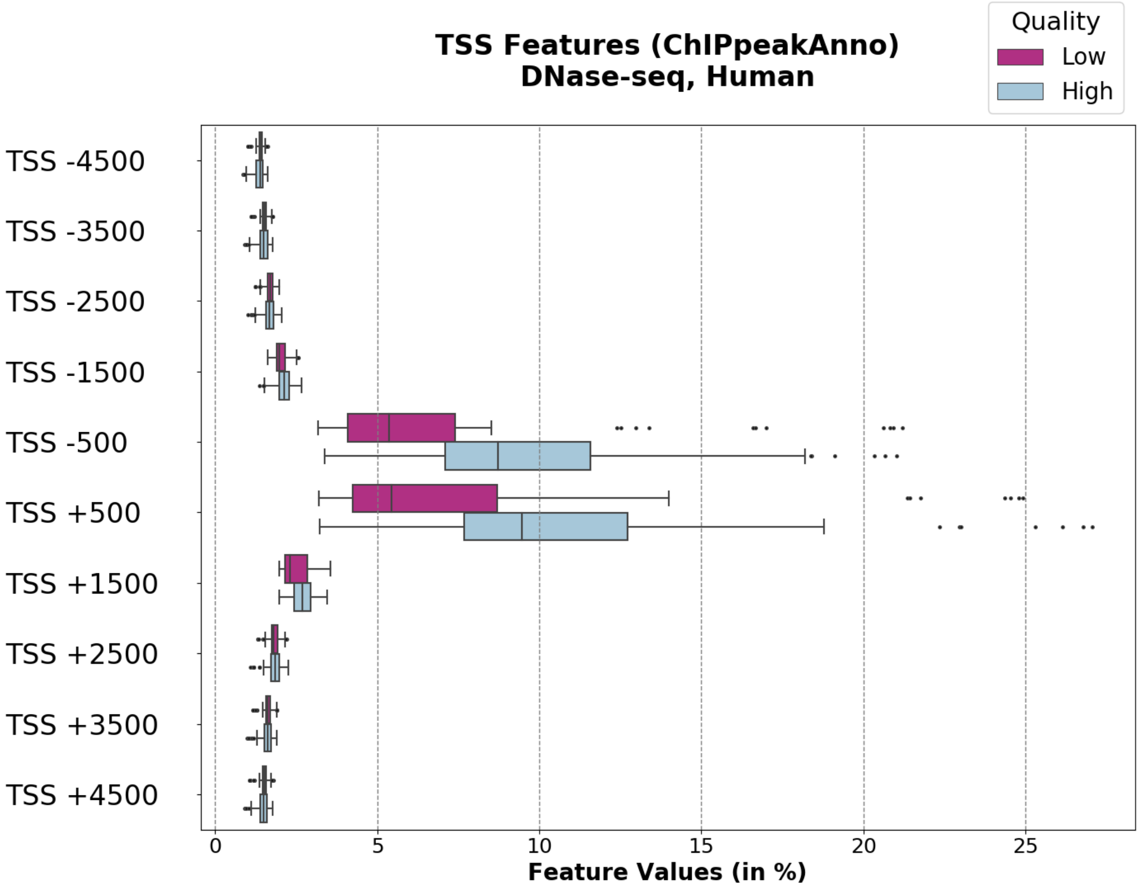
TSS Features (ChIPpeakAnno)
DNase-seq, Human
Quality
Low
High
TSS -4500
TSS -3500
TSS -2500
TSS -1500
TSS -500
TSS +500
TSS +1500
TSS +2500
TSS +3500
TSS +4500
0
5
10
15
20
25
Feature Values (in %)

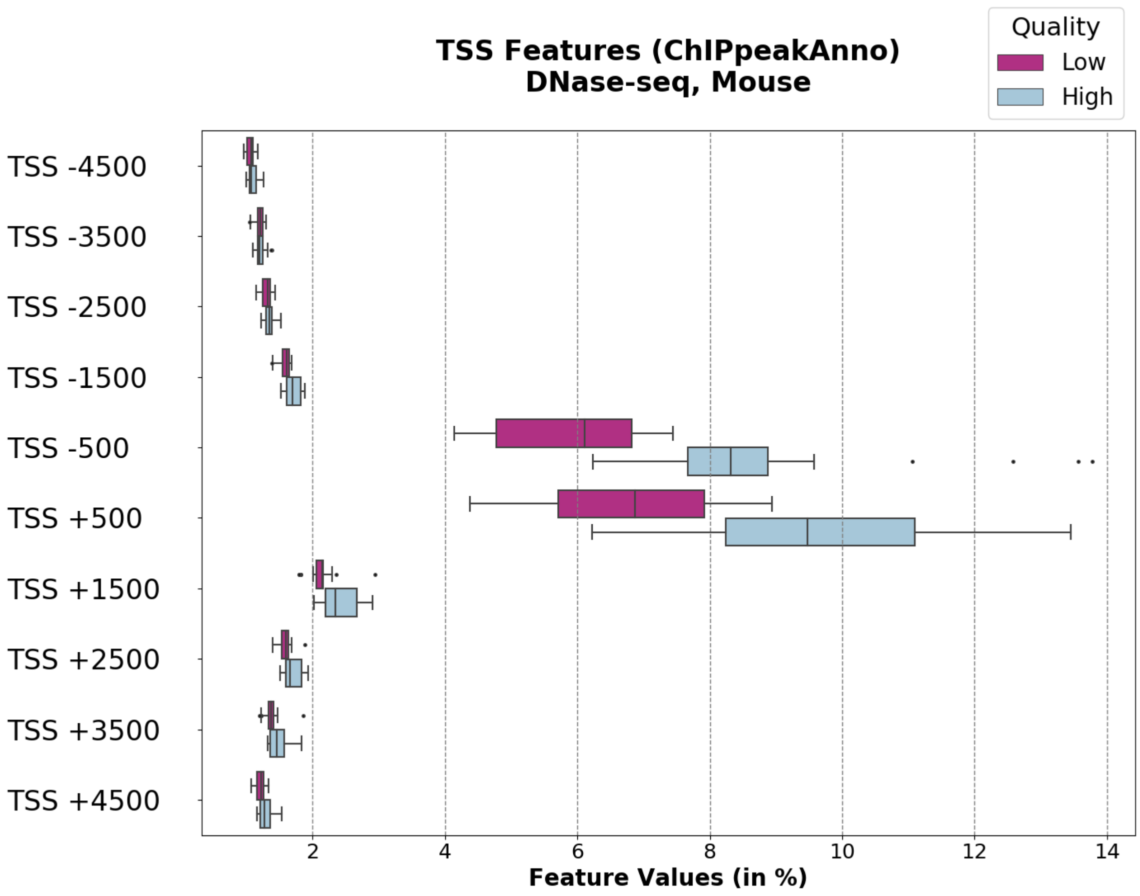

TSS Features (ChIPpeakAnno)
DNase-seq, Mouse
Quality
Low
High
TSS -4500
TSS -3500
TSS -2500
TSS -1500
TSS -500
TSS +500
TSS +1500
TSS +2500
TSS +3500
TSS +4500
2
4
6
8
10
12
14
Feature Values (in %)


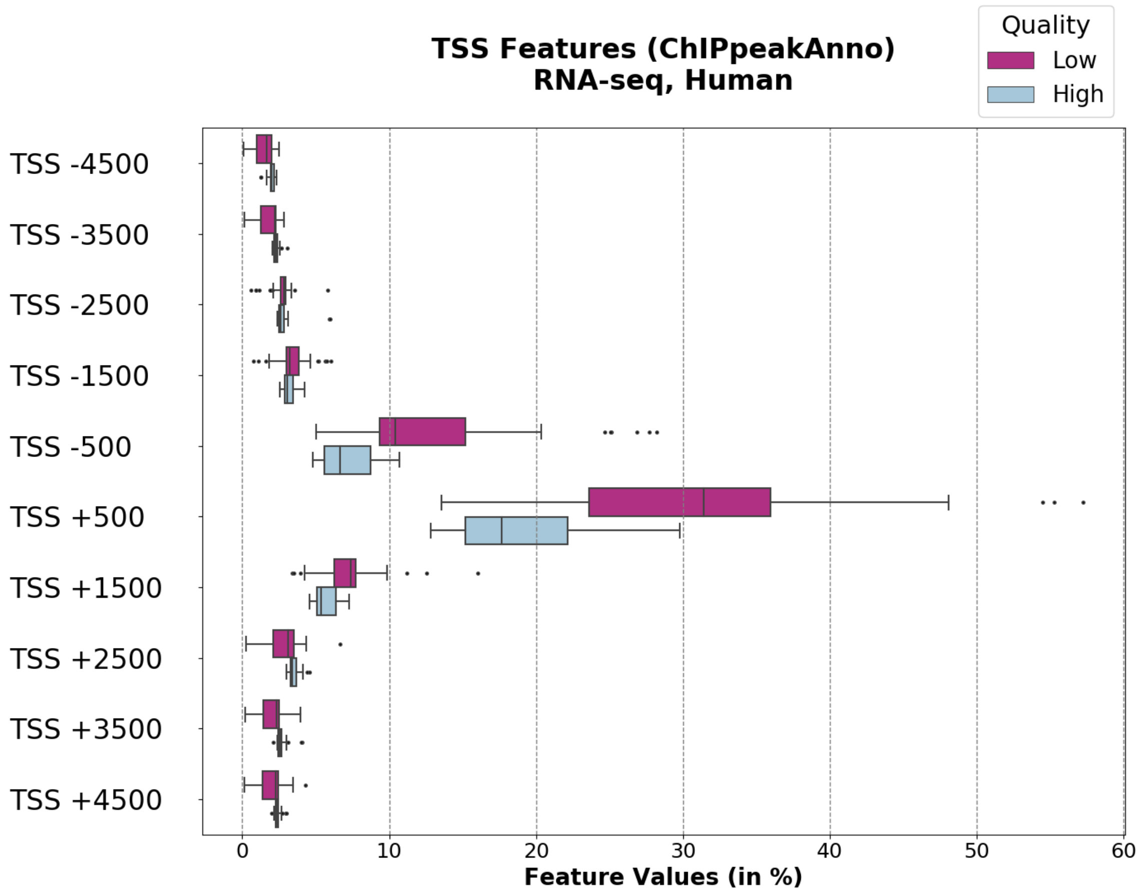

TSS Features (ChIPpeakAnno)
RNA-seq, Human
Quality
Low
High
TSS -4500
TSS -3500
TSS -2500
TSS -1500
TSS -500
TSS +500
TSS +1500
TSS +2500
TSS +3500
TSS +4500
0
10
20
30
40
50
60
Feature Values (in %)

**Figure S2 – Predictive performance of optimal machine learning models**

Models were built for each data subset (y-axis) and all possible combination of feature sets. Several performance measures were used: (**A**) area under ROC-curve (auROC), (**B**) area under precision-recall curve (auPRC), (**C**) accuracy (ACC), and (**D**) F1 measure. Feature sets: RAW (raw data), MAP (genome mapping), LOC (genomic localization), TSS (transcription start sites profile).

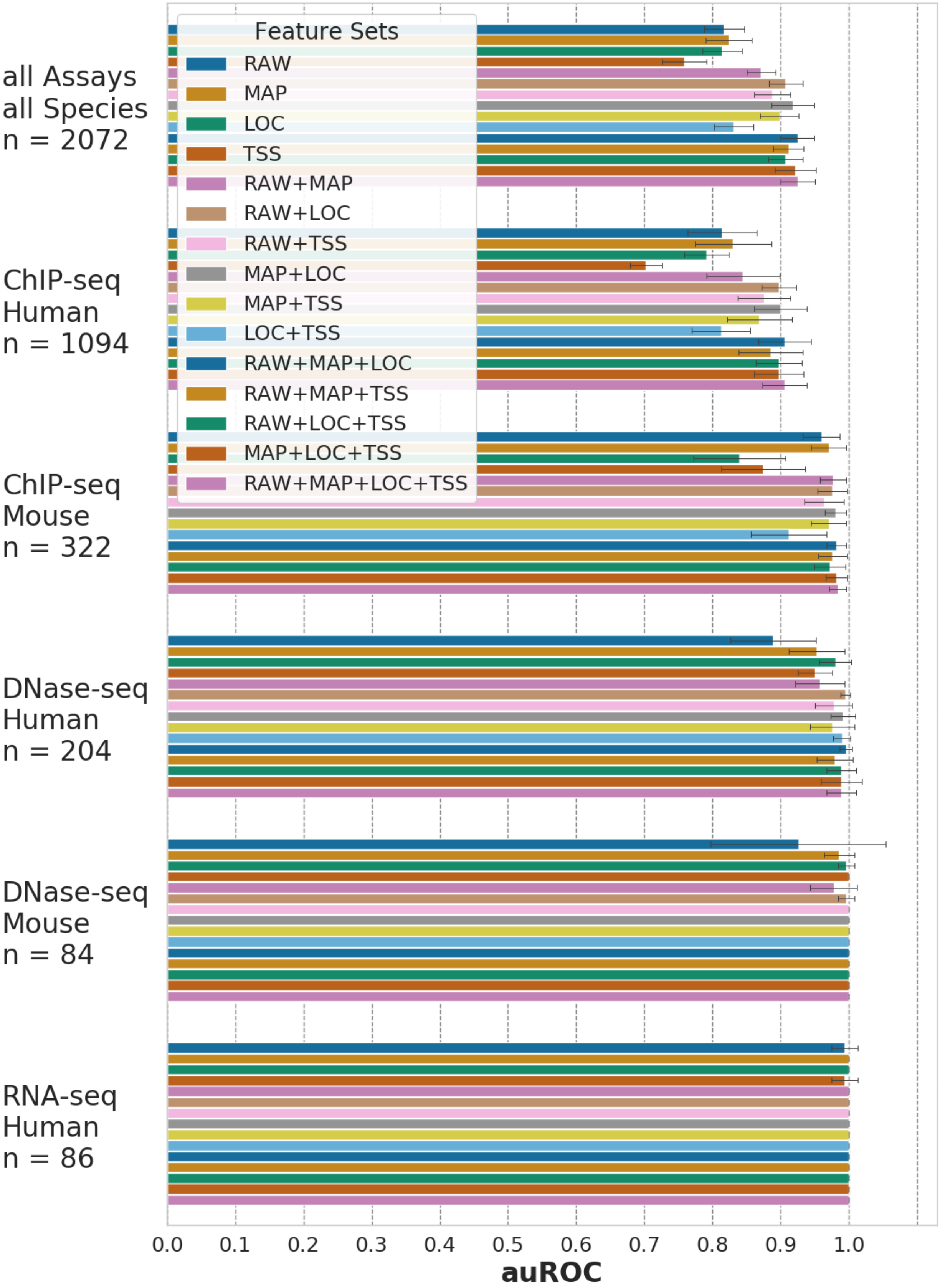
Feature Sets
RAW
MAP
LOC
TSS
RAW+MAP
RAW+LOC
RAW+TSS
MAP+LOC
MAP+TSS
LOC+TSS
RAW+MAP+LOC
RAW+MAP+TSS
RAW+LOC+TSS
MAP+LOC+TSS
RAW+MAP+LOC+TSS
all Assays
all Species
n = 2072
ChIP-seq
Human
n = 1094
ChIP-seq
Mouse
n = 322
DNase-seq
Human
n = 204
DNase-seq
Mouse
n = 84
RNA-seq
Human
n = 86
0.0
0.1
0.2
0.3
0.4
0.5
0.6
0.7
0.8
0.9
1.0
auROC

**A**

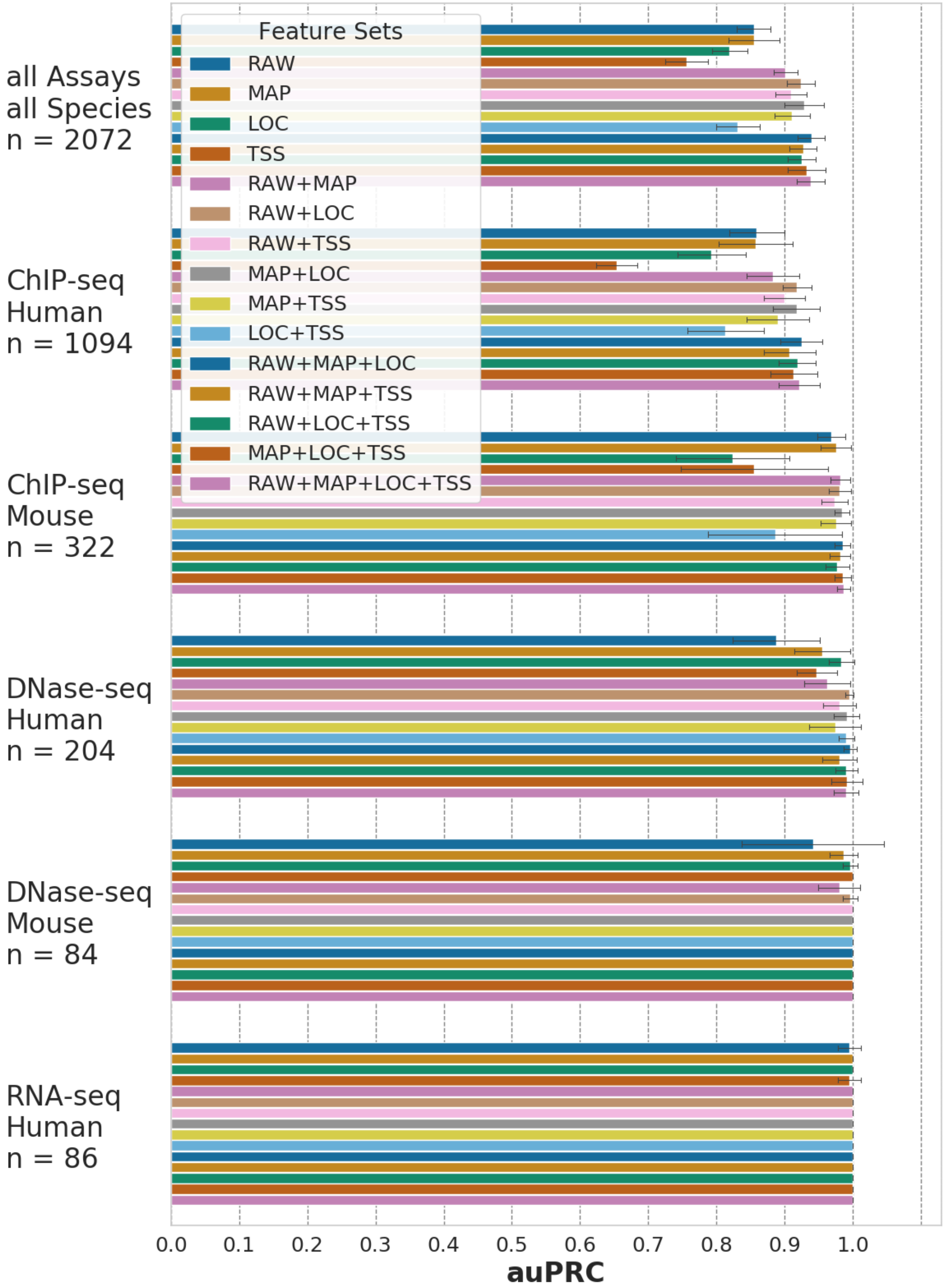

Feature Sets
RAW
MAP
LOC
TSS
RAW+MAP
RAW+LOC
RAW+TSS
MAP+LOC
MAP+TSS
LOC+TSS
RAW+MAP+LOC
RAW+MAP+TSS
RAW+LOC+TSS
MAP+LOC+TSS
RAW+MAP+LOC+TSS
all Assays
all Species
n = 2072
ChIP-seq
Human
n = 1094
ChIP-seq
Mouse
n = 322
DNase-seq
Human
n = 204
DNase-seq
Mouse
n = 84
RNA-seq
Human
n = 86
0.0
0.1
0.2
0.3
0.4
0.5
0.6
0.7
0.8
0.9
1.0
auPRC


**B**

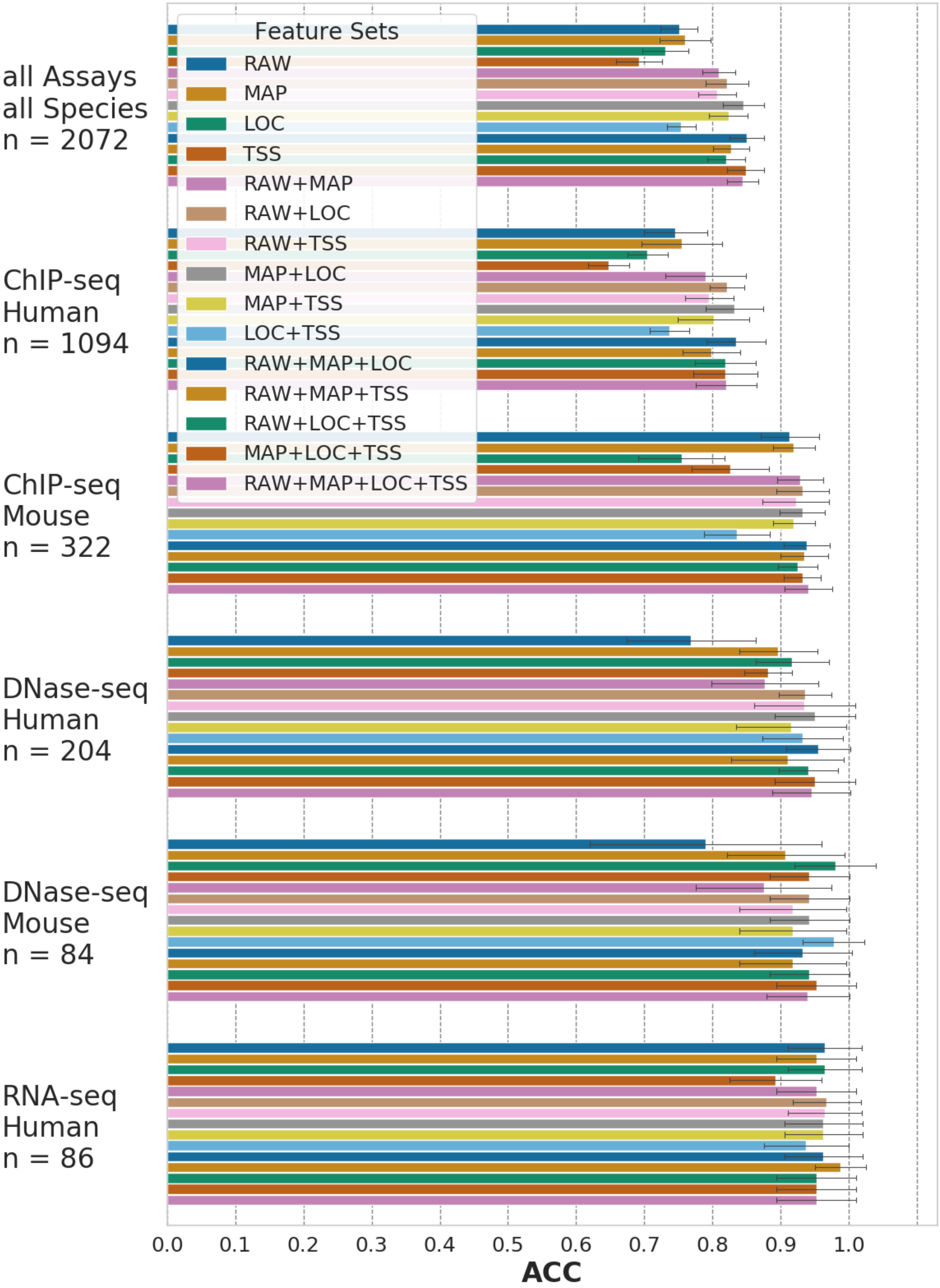
Feature Sets
RAW
MAP
LOC
TSS
RAW+MAP
RAW+LOC
RAW+TSS
MAP+LOC
MAP+TSS
LOC+TSS
RAW+MAP+LOC
RAW+MAP+TSS
RAW+LOC+TSS
MAP+LOC+TSS
RAW+MAP+LOC+TSS
all Assays
all Species
n = 2072
ChIP-seq
Human
n = 1094
ChIP-seq
Mouse
n = 322
DNase-seq
Human
n = 204
DNase-seq
Mouse
n = 84
RNA-seq
Human
n = 86
0.0 0.1 0.2 0.3 0.4 0.5 0.6 0.7 0.8 0.9 1.0
ACC

**C**

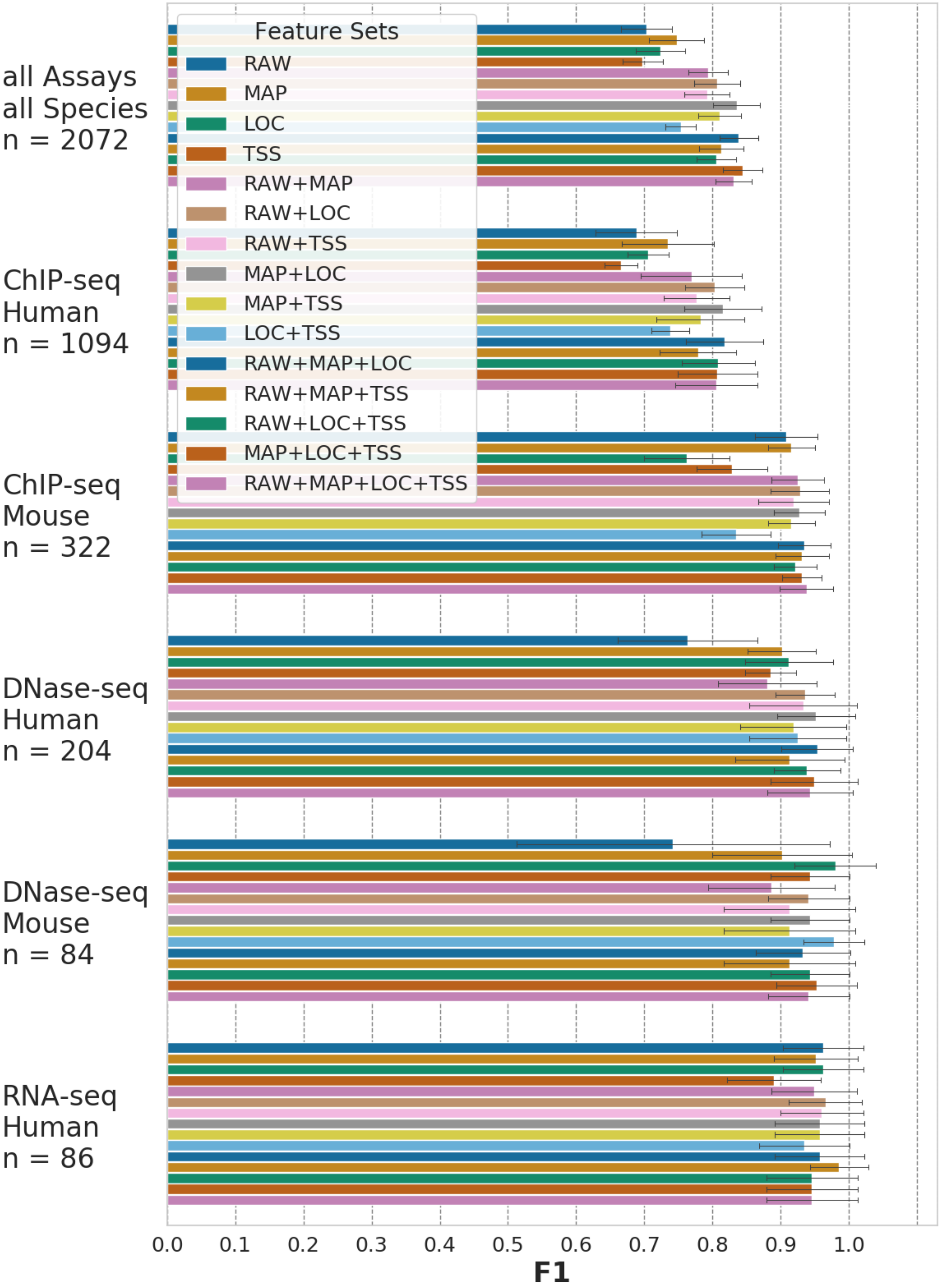

Feature Sets
RAW
MAP
LOC
TSS
RAW+MAP
RAW+LOC
RAW+TSS
MAP+LOC
MAP+TSS
LOC+TSS
RAW+MAP+LOC
RAW+MAP+TSS
RAW+LOC+TSS
MAP+LOC+TSS
RAW+MAP+LOC+TSS
all Assays
all Species
n = 2072
ChIP-seq
Human
n = 1094
ChIP-seq
Mouse
n = 322
DNase-seq
Human
n = 204
DNase-seq
Mouse
n = 84
RNA-seq
Human
n = 86
0.0
0.1
0.2
0.3
0.4
0.5
0.6
0.7
0.8
0.9
1.0
F1


**D**

**Figure S3 – Within-experiment benchmarks**

ENCODE experiment IDs are shown along the y-axis while the files that belong to one experiment are plotted within the area indicated by the dashed lines. The x-axis shows the predictive probability of a file to be of low-quality $P_{low}$.

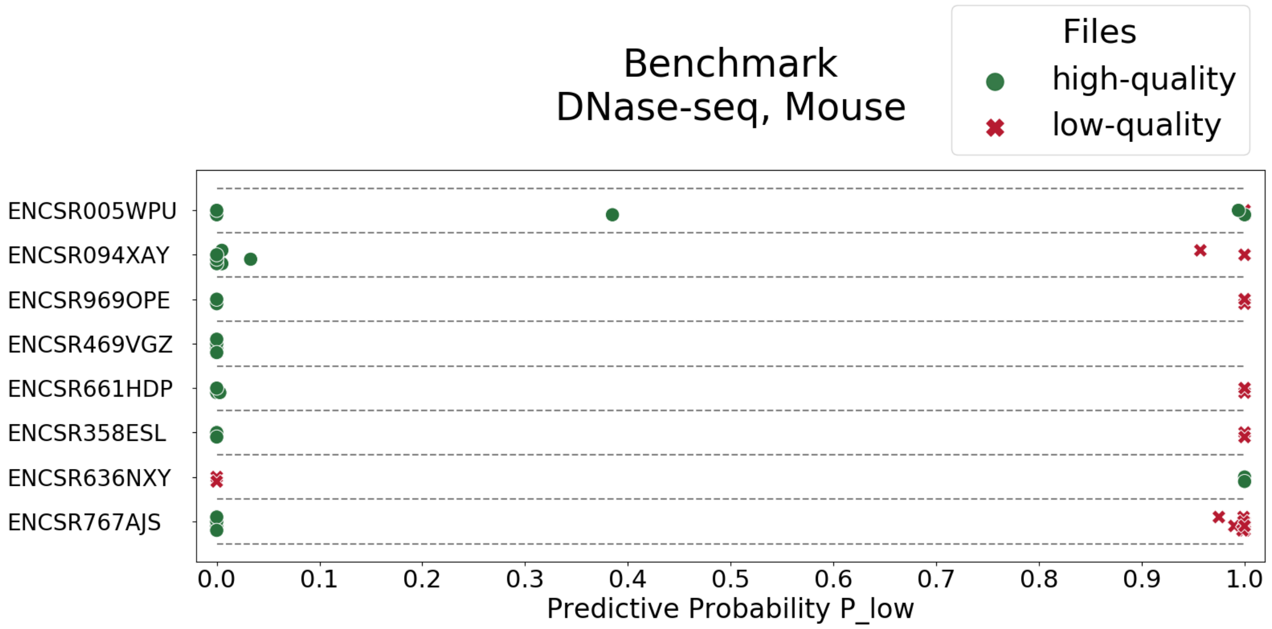


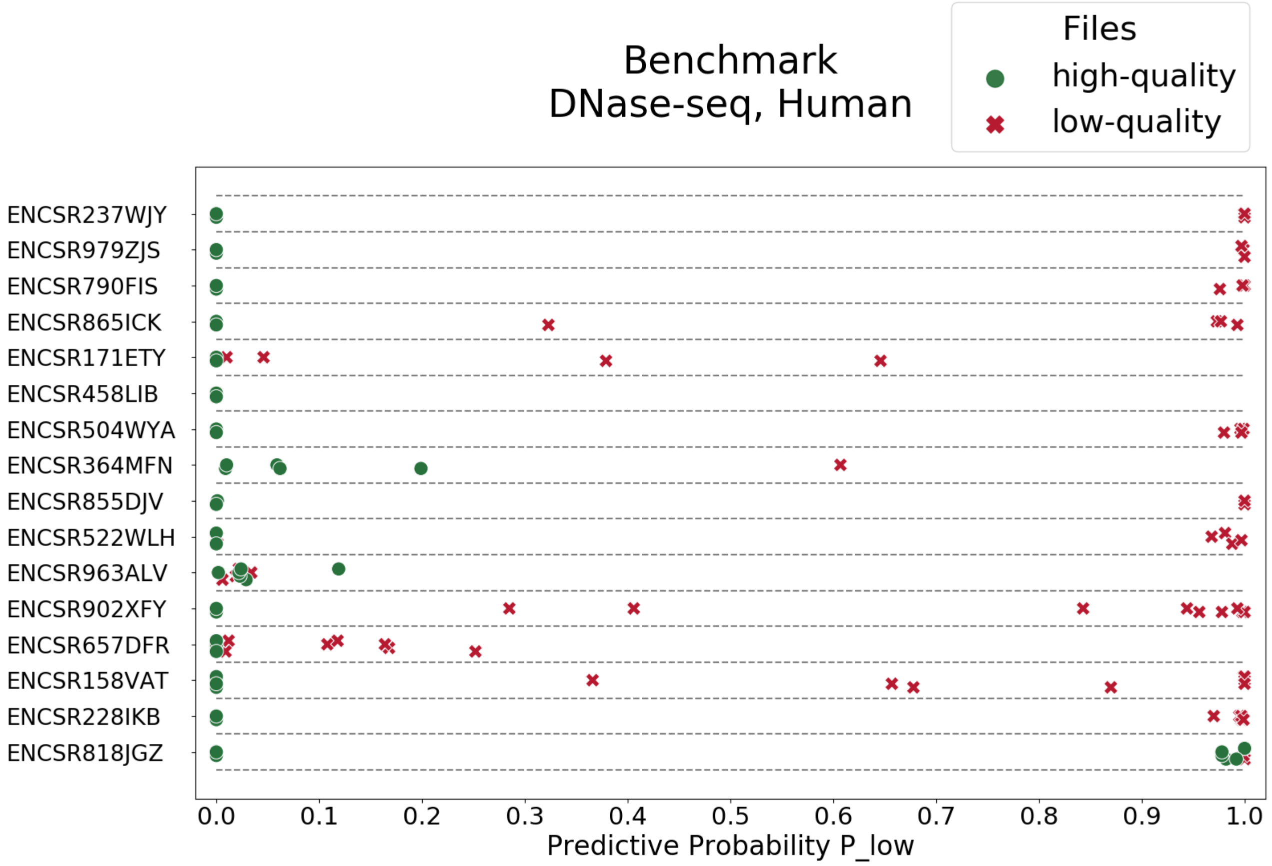

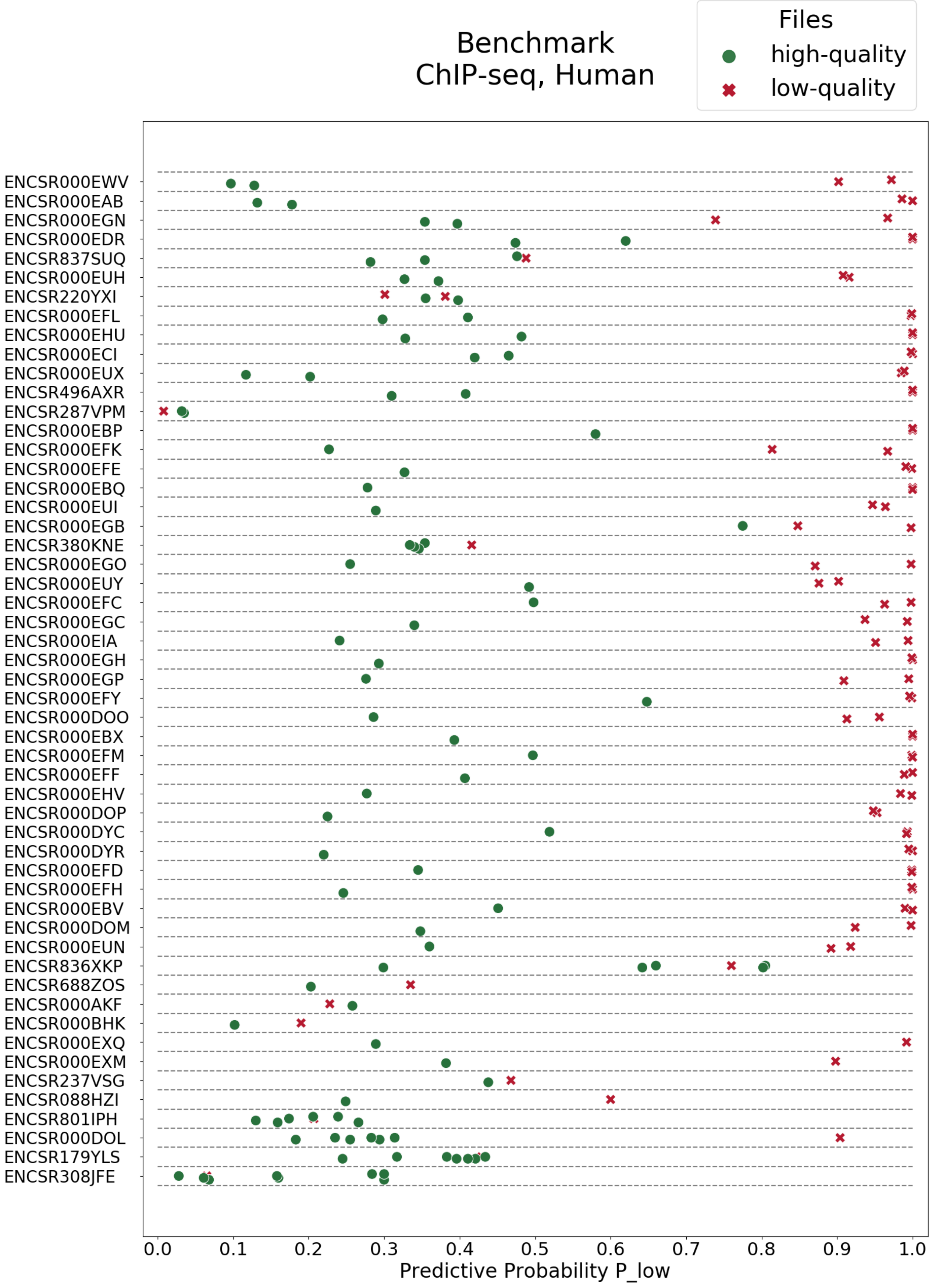

Benchmark
ChIP-seq, Human
Files
high-quality
low-quality
ENCSR000EWV
ENCSR000EAB
ENCSR000EGN
ENCSR000EDR
ENCSR837SUQ
ENCSR000EUH
ENCSR220YXI
ENCSR000EFL
ENCSR000EHU
ENCSR000ECI
ENCSR000EUX
ENCSR496AXR
ENCSR287VPM
ENCSR000EBP
ENCSR000EFK
ENCSR000EFE
ENCSR000EBQ
ENCSR000EUI
ENCSR000EGB
ENCSR380KNE
ENCSR000EGO
ENCSR000EUY
ENCSR000EFC
ENCSR000EGC
ENCSR000EIA
ENCSR000EGH
ENCSR000EGP
ENCSR000EFY
ENCSR000DOO
ENCSR000EBX
ENCSR000EFM
ENCSR000EFF
ENCSR000EHV
ENCSR000DOP
ENCSR000DYC
ENCSR000DYR
ENCSR000EFD
ENCSR000EFH
ENCSR000EBV
ENCSR000DOM
ENCSR000EUN
ENCSR836XKP
ENCSR688ZOS
ENCSR000AKF
ENCSR000BHK
ENCSR000EXQ
ENCSR000EXM
ENCSR237VSG
ENCSR088HZI
ENCSR801IPH
ENCSR000DOL
ENCSR179YLS
ENCSR308JFE
0.0
0.1
0.2
0.3
0.4
0.5
0.6
0.7
0.8
0.9
1.0
Predictive Probability P_low

**Figure S4 – Cross-species generalization**

Good performances of species-specific models in cross-species predictions demonstrate model generalization to other species. ROC curves show the classification performance for different species-assay combinations. The solid lines represent cases in which data from the same species was used to define training and testing sets. The dashed lines show the performance on cases in which the species defining the training data differs from the species defining the testing data; e.g. ”HS → MM” means the training set contains only data from human (Homo sapiens), while the testing data is only from mouse (Mus musculus). Legends also show values of areas under receiver operating characteristics. Feature sets: RAW (raw data), MAP (genome mapping), LOC (genomic localization), TSS (transcription start sites profile).

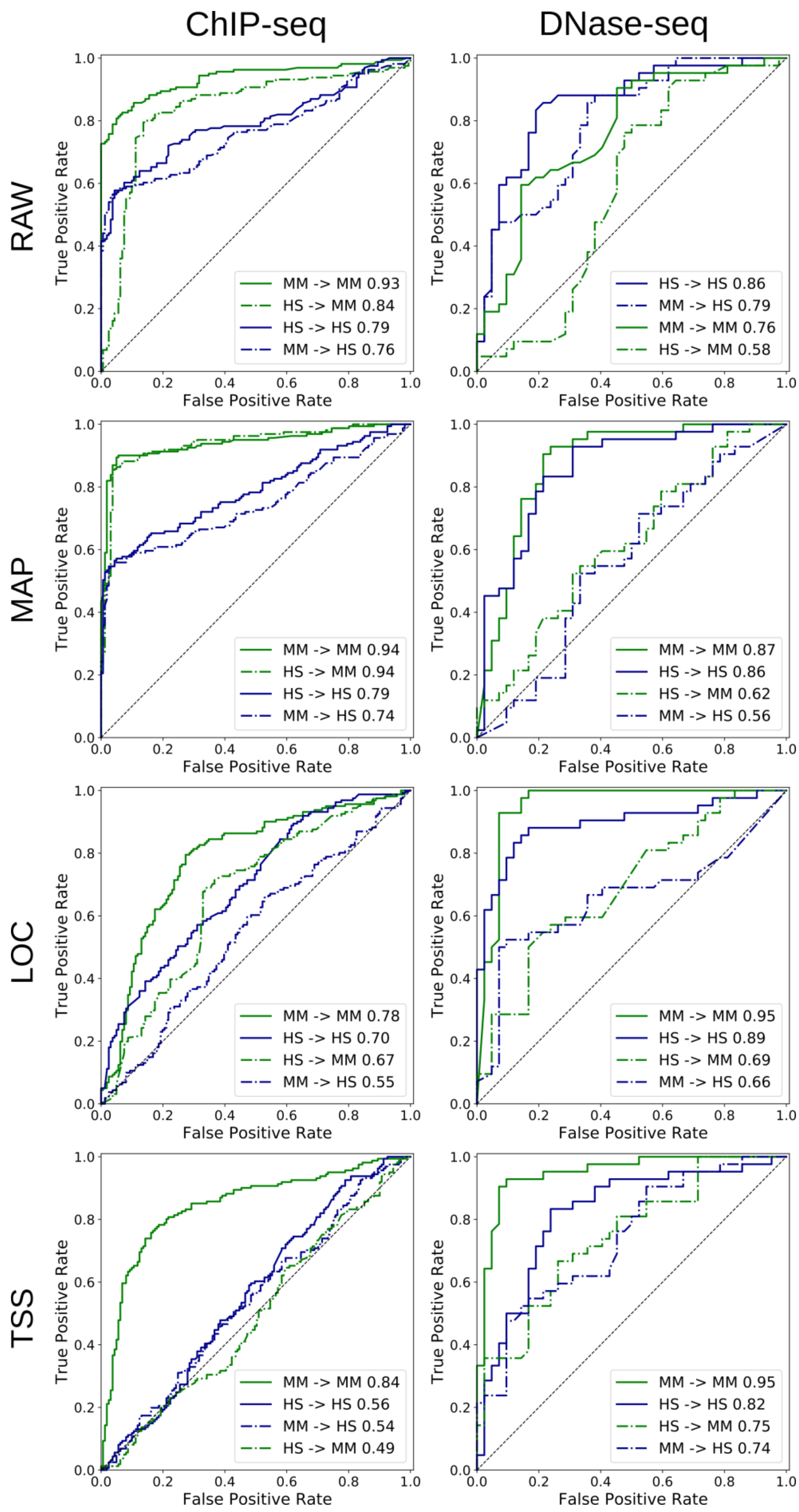
ChIP-seq
DNase-seq
RAW
MAP
LOC
TSS
True Positive Rate
False Positive Rate
MM -> MM 0.93
HS -> MM 0.84
HS -> HS 0.79
MM -> HS 0.76
HS -> HS 0.86
MM -> HS 0.79
MM -> MM 0.76
HS -> MM 0.58
MM -> MM 0.94
HS -> MM 0.94
HS -> HS 0.79
MM -> HS 0.74
MM -> MM 0.87
HS -> HS 0.86
HS -> MM 0.62
MM -> HS 0.56
MM -> MM 0.78
HS -> HS 0.70
HS -> MM 0.67
MM -> HS 0.55
MM -> MM 0.95
HS -> HS 0.89
HS -> MM 0.69
MM -> HS 0.66
MM -> MM 0.84
HS -> HS 0.56
MM -> HS 0.54
HS -> MM 0.49
MM -> MM 0.95
HS -> HS 0.82
HS -> MM 0.75
MM -> HS 0.74

**Figure S5 – Paired-end human ChIP-seq data subset**

**A** Specialized models trained on the human ChIP-seq data subset using different feature sets (y-axis) outperform one-feature predictions except with RAW features. **B** Correlation of predictive performance of a generic model compared to different specialized models demonstrates lack of bias. For each feature set, performance of the generic model (trained on all data subsets) is shown for the human ChIP-seq data subset (green bars) and compared to specialized models trained for the subset only (blue bars). Error bars show standard deviations derived from 10-fold cross-validations within the grid search. Feature sets: RAW (raw data), MAP (genome mapping), LOC (genomic localization), TSS (transcription start sites profile), ALL (all features).

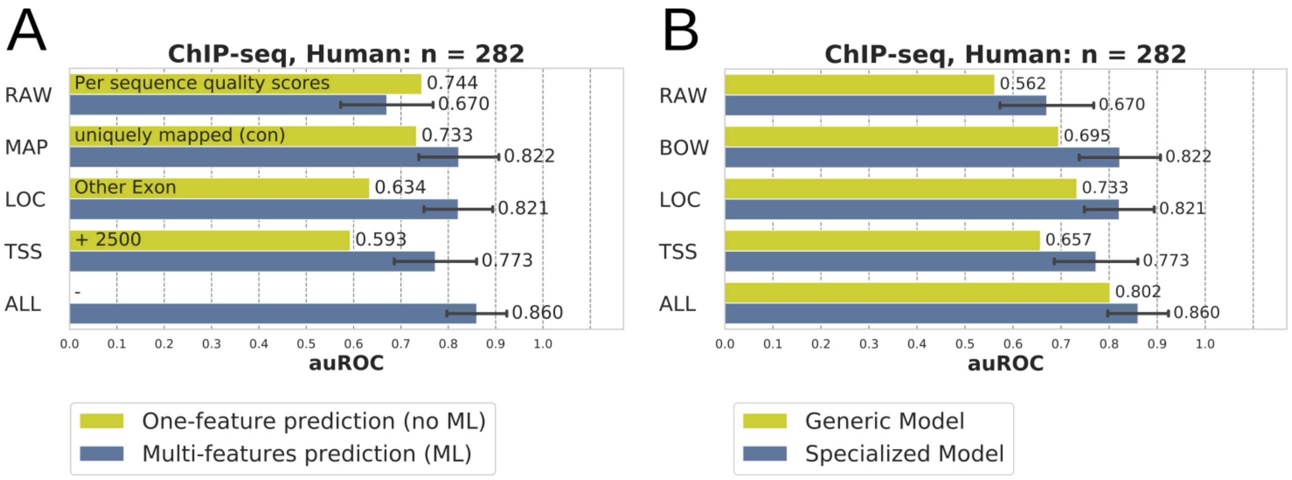
